# Cytokine expression patterns predict suppression of vulnerable neural circuits in a mouse model of Alzheimer’s disease

**DOI:** 10.1101/2024.03.17.585383

**Authors:** Dennis Chun Yin Chan, ChaeMin Kim, Rachel Y. Kang, Madison K. Kuhn, Samuel Cramer, Hayreddin Ünsal, Lynne M. Beidler, Kevin L. Turner, Thomas Neuberger, Nanyin Zhang, Elizabeth A. Proctor

## Abstract

Alzheimer’s disease is a neurodegenerative disorder characterized by progressive amyloid plaque accumulation, tau tangle formation, neuroimmune dysregulation, synapse and neuron loss, and changes in neural circuit activation that lead to cognitive decline and dementia. Early molecular and cellular disease-instigating events occur 20 or more years prior to presentation of symptoms, making them difficult to study, and for many years amyloid-β, the aggregating peptide seeding amyloid plaques, was thought to be the toxic factor responsible for cognitive deficit. However, strategies targeting amyloid-β aggregation and deposition have largely failed to produce safe and effective therapies, and amyloid plaque levels poorly correlate with cognitive outcomes. However, a role still exists for amyloid-β in the variation in an individual’s immune response to early, soluble forms of aggregates, and the downstream consequences of this immune response for aberrant cellular behaviors and creation of a detrimental tissue environment that harms neuron health and causes changes in neural circuit activation. Here, we perform functional magnetic resonance imaging of awake, unanesthetized Alzheimer’s disease mice to map changes in functional connectivity over the course of disease progression, in comparison to wild-type littermates. In these same individual animals, we spatiotemporally profile the immune milieu by measuring cytokines, chemokines, and growth factors across various brain regions and over the course of disease progression from pre-pathology through established cognitive deficit. We identify specific signatures of immune activation predicting hyperactivity followed by suppression of intra- and then inter-regional functional connectivity in multiple disease-relevant brain regions, following the pattern of spread of amyloid pathology.

## Introduction

Alzheimer’s disease (AD) is a neurodegenerative disorder characterized by cognitive impairments that result in a substantial decrease in quality of life metrics (Stites et al 2018, Pan et al 2015). Classical hallmarks of AD are the development and distribution of neurotoxic amyloid-β peptides and plaques (Thal et al 2002, Braak et al 2011, Hampel et al 2021), followed by an accumulation of neurofibrillary tau tangles (DeTure and Dickson 2019, Dickson 1997) and, ultimately, neuronal loss and cognitive impairment (Knopman et al 2021, Kim et al 2022). While modern technologies have led to a vast body of literature describing molecular and cellular pathologies of AD, as well as an understanding of the effect of the disease on global brain function and neural circuitry, the mechanisms by which these molecular and cellular changes result in altered neural circuitry, even before substantial neuron death, remain unexplored. This gap in our understanding has led to a lack of therapeutic strategies capable of halting, reversing, or even substantially delaying the cognitive decline and dementia that are the primary patient concern in AD (Yiannopoulou et al 2019, Venugopalan et al 2021). Here, we make the first steps toward connecting molecular and cellular changes occurring as a result of amyloid-β deposition to specific changes in global brain connectivity and neural circuit alterations in a mouse model of AD.

Healthy cognitive function is enabled by coordinated neural activity in a large-scale brain network, with each aspect of cognitive function having a corresponding physical manifestation in brain activity. Altered activity of a particular neural circuit or network leads to impairment of the associated cognitive function. In both humans and animals, the coordination of neural activity among regions of the brain is predominantly studied using functional magnetic resonance imaging (fMRI), which leverages the magnetic properties of oxygenated blood to quantify which regions of the brain are consistently active (as determined by increased blood flow) at the same time. This correlated activity creates large-scale brain networks that represent brain activity while at rest or accomplishing a specific task. Changes in resting brain networks are an early indicator of cognitive impairment in various neurological disorders, including AD (Filippini et al 2009, Greicius et al 2003, Dai et al 2020, Pievani et al 2014, Choi et al 2021, Yamamoto et al 2019). Even in cognitively normal humans, changes in resting state functional connectivity networks coincide with the early-stage accumulation of amyloid-β, with some subjects later developing cognitive impairment progressing to AD dementia (Palmqvist et al 2017, Moffat et al 2022). Thus, resting state functional connectivity networks, as measured by MRI, are a quantitative and sensitive measure of the neural activity and circuitry that are the physiological manifestation of cognitive function.

The specific molecular and cellular interactions that enable the highly coordinated neural circuit activity underlying cognitive function are highly complex and incompletely understood. The accumulation of amyloid-β in AD pathology evokes a neuroinflammatory response (Dhawan et al 2012, Nordengen et al 2019, Kelley and Petersen 2007). Neurons in distress secrete chemokines that cause microglia, the innate immune cells of the brain, to migrate toward and phagocytize amyloid-β aggregates in an attempt to clear them. Microglia surrounding amyloid plaques are activated, and futile attempts to clear highly stable amyloid plaques evoke a phenotypic shift to become disease-associated microglia (DAM). DAM are dysfunctional in that they no longer fulfill neither their homeostatic nor their phagocytic role in the brain, but express high levels of cytokines (Yan et al 1996, Simard et al 2006, Pan et al 2011), leading to a chronic neuroinflammatory environment (Kinney et al 2018, Wang et al 2023). Neuroinflammation has detrimental effects on neurophysiology, such as increasing the intrinsic excitability of neurons, leading to higher spontaneous firing rates and desynchronized neuro-electrical behavior (Akiyoshi et al 2018, Ren et al 2020, Nguyen et al 2020). Cytokines, the secreted immune cues by which all cells communicate, stimulate intracellular signaling pathways such JAK/STAT and MAPT (Hu et al 2021, Zhang and Liu 2002) that control neuronal fate and function. Cytokine cues also have a profound effect on synaptic plasticity (Nguyen et al 2020, Liu et al 2012, Jung et al 2023), suggesting that microglial immune activity affects neuronal activity and, at a larger scale, potentially alters functional connectivity at the regional and whole-brain levels, and thereby cognitive performance (Nguyen et al 2020, Yamamoto et al 2019).

Defining the multi-scale mechanisms by which neuroimmune interactions regulate neural circuit activity, and how dysregulation of these interactions disrupts coordinated brain functions to result in cognitive impairment and dementia, would allow for the design of therapeutic strategies directly targeting the primary patient concern in Alzheimer’s disease. While there is some understanding of the neural circuits that are selectively vulnerable to Alzheimer’s disease, identification of disease in patients occurs only following onset of cognitive symptoms, which is late in the timescale of the disease-driving molecular and cellular events that begin as early as 20 years before cognitive impairment. While these presymptomatic disease-driving events, including dysregulation of immune interactions, can be (and are) studied in animal models, the study of global brain function and neural circuitry in model animals is complicated by the need for them to remain both still and conscious to undergo fMRI. To address this gap, we have developed an awake-rodent paradigm for functional MR imaging and adjusted it for the unique needs of animals with dementia. Here, we investigate the evolving relationship between immune interactions and neural circuit function in a neurodegenerative disease context using a well-established animal model of AD-like amyloid-β accumulation: the 5xFAD mouse. We profiled the molecular immune environment with regional specificity and quantified the corresponding functional connectivity over the course of disease development, and found that brain regions experiencing amyloid-β accumulation experience a breakdown in intra-regional functional connectivity preceding the loss of communication between that region and the rest of the brain. These losses correspond to the appearance of a signature of cytokines and chemokines indicating microglial chemotaxis and activation, spreading spatiotemporally through the brain in the same pattern as amyloid deposition, suggesting amyloid-driven microglial involvement in the suppression of neural circuitry responsible for cognitive impairment in AD.

## Methods

### Animal subjects

5xFAD hemizygous B6SJL (MMRRC Strain #034840-JAX) background strain and wildtype mice were used to breed a colony of mice that were either hemizygous for the 5xFAD related genes, or wild-type littermates. All mice were genotyped following the protocol provided by Jackson Labs (Protocol Number: 31769). All mice were also genotyped for the retinal degenerative gene Pde6B using the protocol provided by Jackson Labs (Protocol Number: 31378). Animals that were found to be homozygous for the Pde6B gene were excluded from the study. Both male and female animals were included in the study (animal metadata details provided in **Supplementary Table 1**). Mice were singly housed following the headpost surgery (procedure below) at 1 month old. Food and water were provided ad libitum. Nesting material and chew blocks were provided to singly housed mice as enrichment. The housing room adhered to a 12-hour light and 12-hour dark cycle. The study consisted of 158 mice: 27 for longitudinal imaging studies (had imaging done in all four time points of 1.5, 2, 4 and 6 months), and the rest are used for cross-sectional experiments at single time points. Both groups (longitudinal and single) were included in the imaging experiments, with each timepoint from a longitudinal animal treated as an individual subject. Thus, the experiment featured 31 mice at age 1.5 months, 35 at age 2 months, 32 at age 4 months, and 33 at age 6 months. All mice underwent rs-fMRI. Each brain region’s cytokine protein levels were profiled in 105 animal brains from the same cohort of animals that underwent rs-fMRI. The breakdown of animals per group is region dependent and are provided in **Supplementary Table 2**. All experiments were conducted during the dark cycle to better capture the properties of the awake state, with euthanasia and tissue collection (procedure described in ‘Brain Extraction’) executed a week after imaging. All experiments in the present study were approved by the Institutional Animal Care and Use Committee (IACUC) at the Pennsylvania State University (PRAMS201647005).

### Surgical Procedures

Mice had headpost implantation surgery done at 30 days old. Mice were anesthetized with 3% isoflurane mixed with oxygen at the flow rate of 1.0 L/min in an isoflurane chamber. The surgical plane of anesthesia was confirmed by pinching the lower limbs of the mouse. The scalp of the mouse was shaved using an electric clipper. A heating mat with a rectal probe (PhysioSuite, Kent Scientific) was used to maintain a body temperature around 36-37 °C. A custom surgical setup that replicates the awake animal restrainer used in neuroimaging experiments is utilized for headpost surgery. Once the mouse was fixed in the surgical frame, the scalp was cleaned twice alternating between provodine iodine and 70% isopropyl alcohol. Ophthalmic ointment was applied to the animal’s eyes to prevent dryness during the course of surgery. The scalp was infiltrated with Bupivacaine (4mg/kg), followed by an anterior to posterior incision before excising the skin to form a circular excision anterior to the ears, and posterior to the eyes. All relevant soft tissue was removed, leaving only the skull exposed. The site was washed one time with sterilized saline solution. Once dried, an etching solution (composed of citric acid, C&B Metabond) was applied to the skull for 30 seconds before being rinsed off using saline solution. Dental cement (C&B Metabond) consisting of a ratio of 1 drop catalyst (C&B Metabond), 4 drops of “quick!” base (C&B Metabond), and 2 scoops of clear L powder (C&B Metabond) was applied to the exposed skull. Once set and cured, a custom head post constructed from polylactic acid (PLA) was adhered to the dental cap using dental cement of the aforementioned recipe. Upon completion of the procedure, meloxicam (5 mg/kg) was administered 2 days post-surgery. After successful recovery based on home cage behavior and the magnitude of weight change post-surgery, the mouse was moved to subsequent experiments detailed below.

### Acclimation Procedure

Mice were acclimated to the restrainer and MRI scanning environment for awake animal imaging for four days. An acclimation box was used alongside custom-made holders that mimicked the interior of a scanner. All mice were first anesthetized using 3% isoflurane, and an oxygen flow of 1.0 L/min. Once anesthetized, mice had their front limbs bounded together using tape before restraining the mice by attaching a custom-made head bar that connected the implanted headpost to the custom-made restrainer. Once the animals were confirmed to be awake, they were given five minutes to adjust to the restrainer before the acclimation process. The first day of acclimation was 15 minutes with no sound. The second, third and fourth day of acclimation was 30 minutes, 45 minutes, and 60 minutes respectively. A soundtrack that had the recording of the noise during an fMRI session was played for the stated durations at the second, third and fourth day at 110 db.

### fMRI Imaging

rsfMRI was conducted in animals at the awake state on a 7T MRI system interfaced with a Bruker Console (Billerica, MA) housed at the High Field MRI Facility at Pennsylvania State University. T2*-weighted images were acquired using a gradient-echo echo-planar imaging (EPI) sequence with the following parameters: repetition time (TR) = 1.5 second, echo time (TE) = 15 milliseconds, flip angle = 60°, matrix size = 64 x 64, FOV = 1.6 x 1.6 cm^2^ number of slices = 16, slice dimensions = 0.25 mm x 0.25 mm x 0.75 mm. Each EPI acquisition started with 10 or 32 dummy volumes to ensure the MRI signal is at a steady state before data acquisition. A trigger delay of 10 milliseconds was set at the beginning of each volume acquisition such that real time monitoring of respiration rate can be synchronized to the volume acquisition. Respiration was monitored using a custom-made nose cone perforated with angled holes to allow carbon dioxide to diffuse out and oxygen to diffuse in without significant loss of pressure as seen in Supplementary Figure 1A. The respiration monitoring system was connected to the 7T MRI system using a MR-compatible monitoring and gating system (Model 1030; Small Animal Instruments Inc., Stony Brook, NY, USA), which was also interfaced with a desktop that housed the respiration monitoring software from Small Animal Instruments Inc. A total of 4 or 5 EPI scans were collected for every imaging session. Following the completion of EPI acquisition, a structural T1-RARE image was collected. The imaging parameters are as follows: TR = 1500 milliseconds, TE = 8 milliseconds, flip angle = 90°, matrix size = 256 x 256, FOV = 1.6 x 1.6 cm^2^, number of slices = 16, slice dimensions = 0.25 mm x 0.25 mm x 0.75 mm.

### fMRI Image Preprocessing

fMRI images and the corresponding T1-RARE structural images were preprocessed following a paradigm adapted from an in-house pipeline using MATLAB 2022b (Dopfel et al 2019, Liu et al 2020). Briefly, framewise displacement (FD) of individual rsfMRI volumes was determined using the following equation:

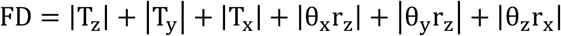

in which T_x_ is the translation in the x axis, T_y_ is the translation in the y axis, T_z_ is the translation in the z axis, θ_x_ is the angle of rotation about the x axis, θ_y_ is the angle of rotation about the y axis, θ_z_ is the angle of rotation about the z axis, r_z_ is the radius of a sphere representing the brain size in the z axis, and r_x_ is the radius of sphere representing the brain size in the x axis. The radius across all 3 dimensions is 5 mm.

rsfMRI frames with displacement that exceeded half a voxel size in the x/y axis (0.125 mm) were discarded, along with one volume preceding it, and one volume after it. Scans that retained 90% or more volumes after motion removal were kept for subsequent processing. Post motion scrubbing, each scan underwent the process of co-registration and normalization. First, the first frame for each EPI scan is co registered to the relevant subject’s T1-RARE structural image using an in-house GUI. Following co-registration, the subject’s T1RARE image was then normalized to a reference brain atlas based on the Allen Brain Atlas (Allen Reference Atlas – Mouse Brain [brain atlas], Available from atlas.brain-map.org). After successful normalization, the affine transformation matrix from the normalization process was then applied to the co-registered EPI images to align the EPI images to the reference brain atlas. Subsequently, the scan was motion corrected using Statistical Parametric Mapping (SPM12, http://www.fil.ion.ucl.ac.uk/spm). For every motion-corrected EPI scan, a custom mask was manually drawn using the first rsfMRI volume for each individual scan. The custom mask excludes areas with significant signal distortion, and is subsequently applied to all volumes in the same scan. White matter and cerebral spinal fluid (WM-CSF) masks were constructed using the reference atlas (Allen Reference Atlas – Mouse Brain [brain atlas], Available from atlas.brain-map.org). The signal in the WM-CSF mask was then used as a nuisance regressor alongside the motion parameters calculated from the motion correction step in a general linear model. After regression, rsfMRI images were spatially smoothed using a 3x3 Gaussian kernel (FWHM = 0.50 mm) followed by bandpass temporal filtering (0.01 - 0.1 Hz).

### fMRI Image Postprocessing

The whole brain was divided into 51 (listed in **Supplementary Table 3**) regions of interest (ROI). For each scan, the BOLD time course of each ROI was obtained by regionally averaging the preprocessed BOLD time courses of individual voxels within the ROI. Pair-wise functional connectivity between every two ROIs was then calculated using the Pearson correlation coefficient (i.e. r value) between the corresponding ROI time courses, generating a connectivity matrix for the scan. After generating connectivity matrices for every scan, the matrices underwent Fisher z transformation (using the arctanh function) before averaging within subject connectivity matrices to get a subject average. The same process was repeated for group averages as well (i.e. averaging Fisher z-transformed r values per subject and the averaged z score was transformed back to the r value). The initial evaluation of the entire dataset comprised of examining the correlation matrices in a univariate manner, where the Fisher z transformed pairwise correlation values are used. This is followed by implementing a linear mixed effect model of the following structure:

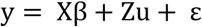

Where y is the response vector with n x 1 dimensions, n being the number of subjects. X is defined as the predictor matrix, with n × p dimensions, where p is the number of fixed effect predictors (dependent on analysis), β is the p × 1 vector of fixed effect coefficients. Z is a n × (q · m) matrix where there are q random effects and m groups. u is defined as a random vector with dimensions q · m × 1 for m groups. ε is the residual vector for n groups. The random effects are the subjects while the fixed effects are a combination of genotype and timepoint, dependent on the statistical question being evaluated (e.g. time point segregated data will only have genotype as the fixed effect to evaluate differences at a single time point). A two-way ANOVA or two-tailed t-test (depending on the model) is conducted to evaluate the statistical significance of the coefficients derived from the linear mixed effect model.

There are two notable functional connectivity metrics referenced in the results: average seed to whole brain functional connectivity; average interregional and intra-regional functional connectivity. Average seed to whole brain connectivity is quantified by taking the calculating the average seed ROI to every other ROI functional connectivity by first z-transforming all of the functional connectivity metrics, taking the average, and then re-transforming back into r space. Average seed ROI to cortical regions is quantified by taking the seed ROI – every other cortical ROI functional connectivity and quantifying the average in the aforementioned manner. Average interregional connectivity is quantified by grouping all the anatomical region’s ROIs and the relevant connectivity’s with other ROIs in different anatomical regions and calculating the mean functional connectivity as mentioned before. Average intraregional connectivity is quantified in the same manner as interregional connectivity with the ROI-ROI connectivity being within the same anatomical system.

### Tissue collection

Mice were euthanized via decapitation by placing the awake mouse into a decapicone and using a guillotine. The decapitated head was then sprayed with 70% sterile ethanol. The skin and muscle are removed from the skull by removing the skin surrounding the cement cap. Forceps are used to grasp the head post attachment and apply a measured amount of force to slowly rock the dental cap back and forth to slowly loosen the cement cap from the skull. Once the cement cap is ‘peeled’ off from the skull, dissection scissors are used to cut up the side of the spinal cord and around the base of the skull at the lateral and ventral borders. The scissors are then inserted at the base of the skull and are used to make a cut down the midline of the skull from posterior to anterior direction. The dorsal part of the skull is then removed and the brain is slowly and carefully dislodged and placed in a separate cell culture dish containing chilled dissection media (x mL of HBSS and x/90 mL 1 M HEPES) hosted within a bed of ice and placed on top of a chilled copper plate.

The brain is then halved into two separate hemispheres, with the right hemisphere to be dissected into individual brain regions. The brain regions are the Occipital Lobe, Temporal Lobe, Parietal Lobe, Frontal Lobe, Hippocampus, Basal Ganglia, Thalamus and Hypothalamus. Each individual brain region is placed into their own respective centrifuge tubes that holds a mixture of lysis buffer and protease inhibitor. The brain regions are homogenized by mechanical titration before sitting for 20 minutes on ice before being centrifuged at 5000 rpm for 5 minutes at 4°C. The supernatant is then placed into a separate tube, and flash frozen in liquid nitrogen and stored at a -80 °C freezer until further use.

### Quantification of protein

Total protein content in the supernatant was quantified using the Pierce BCA Protein Assay kit (Fisher #23225). Manufacturer’s instructions were followed accordingly. Samples were run using technical duplicates and absorbance was quantified using a SpectraMax i3 minimax 300 imaging cytometer (Molecular Devices). Linear regression was used to quantify protein concentration.

### Luminex Multiplexing Immunoassays

Cytokine concentrations of dissected homogenized brain regions from 5xFAD mice were quantified using the Bio-Plex Pro Mouse Cytokines Grp1 Panel 23 Plex (Cat. #M60009RDPD) using the Luminex FLEXMAP3D platform. Manufacturer’s protocol was performed with minor modifications to accommodate the use of a 384 well plate (Kuhn et al 2023), including magnetic beads and antibody solutions used at reduced volume.

### Cytokine data processing and analysis

The Xponent software provided by the Luminex System was used to interpolate sample cytokine concentrations using standard curves derived from using the 5 point logistic regression model. Concentrations below or above the standard limit are either set to 0 pg/mL or the maximum concentration on the curve respectively. An in-house pipeline was used to clean the cytokine data (Kuhn et al 2023): https://github.com/elizabethproctor/Luminex-Data-Cleaning (version 1.05). All cytokines with above-background values for at least half of the subjects were used in further analysis of the cytokine data; we included cytokines with fewer non-zero values if the non-zeros appeared biased toward a particular group. We utilized a linear, supervised multivariate regression model known as Partial Least Squares (PLS) (Wold et al 1984, Wood et al 2015, Kuhn et al 2023, Fleeman et al 2023) to model cytokine signatures of disease and healthy control states. The ROPLS package in R (Thévenot et al 2015) was used to run Projection to Latent Structures/Partial Least Squares Regression (PLSR) or Projection to Latent Structures/Partial Least Squares Discriminant Analysis (PLSDA). PLSR was used to construct a predictive model to model a continuous response variable, such as the timepoint of our covariates. PLSDA models were used to quantify how cytokine signatures could be used to classify diseased brains against healthy brains. All cytokine data was normalized.

Random sub sampling cross validation tests were conducted to select the appropriate number of latent variables (LVs) for PLSR models. Test sets and training sets were determined by randomly sampling the parent dataset. The number of k-folds, consequentially determining the size of the training and test sets, was determined by the following criteria (Kuhn et al 2023): k-Folds = 3 if the number of samples exceeded 30, or k-folds = 5 if the number of samples building model was less than 30. K-fold cross validation was conducted a hundred times. 3 models consisting of either 1, 2, or 3 LVs were constructed in every iteration of the Cross-Validation process. All models were then used to predict the response variable, and the root mean squared error of cross validation (RMSECV). RMSECV has the following formulae:

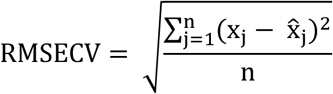

Where x_j_ is the predicted value for the j^th^ sample, 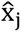 is the actual value for the j^th^ sample, and n is the number of samples in the test set. Every iteration resulted in a completely new random sampling of the parent dataset to produce both the test and training datasets for cross validation. The number of LVs was determined by selecting the models that had the lowest RMSECV, or for PLSDA models the highest predictive accuracy. The subsequent model is then used for subsequent analysis and significance testing.

Significance testing of the model was conducted by permutation testing. 1000 permutations were done to construct a null distribution of random models, where each iteration involved the scrambling of the response variable of interest with respect to the covariates. The final model’s predictive accuracy or RMSERCV is then used to calculate a z score:

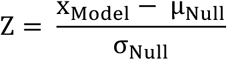

Where X_Model_ is the model’s predictive accuracy or RMSECV, and μ_Null_ is the mean predictive/RMSERCV of the null distribution, and σ_Null_ is the standard deviation of the null distribution simulated by the permutation process. We then calculate the corresponding p value for the significance of the model by comparing the calculated Z score to the Z distribution.

The models are all orthogonalized for improved interpretability. The orthogonalization process results in projecting the maximal amount of covariance in the response vector and the covariate matrix onto the first latent variable (Trygg and Wold 2002). The variable importance in projection (VIP) score was used to threshold which cytokines drove the separation between classes with respect to the response of interest (Galindo-Prieto et al 2014). Thus, loadings from the first latent variable for cytokines that had a VIP score greater than 1 are designated as cytokines of interest, as they are implied to have a greater than average contribution to the model.

### Graph Theoretical analysis

Every functional connectivity matrix (the preprocessed correlation matrix) is converted into a weighted adjacency matrix where the absolute value of the correlation values is used as the weights of each edge. All graph organizational metrics were calculated using the Brain Connectivity Toolbox (Rubinov and Sporns 2010). The graph theory metrics used to determine the hub score were quantified on the group average correlation matrix for every group. The mathematical definitions for the four topological features are described below:

Strength (Sporns 2018, Onnela et al 2005):

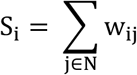

Where S_i_ is defined as the strength for the i^th^ node. W_I,j_ is defined as the weight of the edge between node i and its j^th^ neighbor, and N is the set of all nodes. Strength quantifies the degree of connection between a node and its neighbors.

Clustering Coefficient (Sporns 2018, Onnela et al 2005):

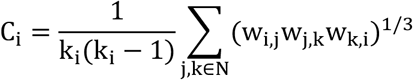

W_i,j_ is the weight between node i and node j, and k_i_ is the number of neighbors of a vertex. The clustering coefficient can be interpreted as the degree to which nodes tend to cluster together, and thus implies the modularity of a network. For the local clustering coefficient, C_i_, the coefficient quantifies how close the neighbors are to node i to being a clique. N is the set of all nodes.

Global Efficiency (Sporns 2018, Latora and Marchiori 2001):

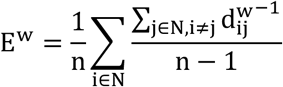

N is the set of all nodes, n is the total number of nodes, and 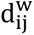 is the weighted shortest path between nodes i and j. Efficiency is essentially the mean of all reciprocals of the weighted distances in a network. The metric quantifies how efficient information is exchanged within a network.

Assortativity (Sporns 2018, Newman 2002):

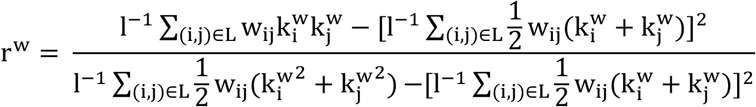

Where 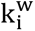 and 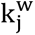 are the weighted degrees of nodes i and j. w_ij_ is the weight of the edge between nodes i and j, l is the total number of edges, While L is the set of all edges within the network. Assortativity is used to determine the degree to which nodes connect to other nodes with similar properties within the network, and can be interpreted as a metric of network resilience.

### Correlation between Neuroinflammation and Functional Connectivity Metrics

Spearman’s rank correlation was conducted to investigate the relationship between cytokine signatures derived from a PLSDA model of Month 6 cytokine concentrations and either interregional system functional connectivity or seed to whole brain functional connectivity. Both the PLSDA model and the functional connectivity metrics have been described in prior sections. Statistical testing in these correlation models were one-tailed, with the alternative hypothesis testing that ρ > 0 and the null hypothesis is ρ ≤ 0.

## Results

In order to investigate the relationship between regional neuroinflammation and functional connectivity of the brain over the development of Alzheimer’s disease, we conducted resting state fMRI and multiplexing experiments of dissected brain regions at four different time points in the 5xFAD rodent model of Alzheimer’s disease. A custom headpost restrainer was used to acquire awake resting state fMRI data (Supplementary Figure 1A). We conducted multiple linear mixed effects models (LME) to understand how disease progression affected the functional connectivity at the brain at the individual and anatomical level. From there, we proceeded to conduct another LME analysis involving metrics from graph theory to understand how disease severity relates to brain network changes at the global level. In parallel to the functional analysis, we also quantified regional neuroinflammatory profiles through the use of linear multivariate models such as projection to latent space (PLS). We observed that both the functional connectivity and neuroinflammatory results indicate that the intermediate stage of amyloid-β pathology is correlated with the greatest degree of separation between the 5xFAD and WT groups. Additionally, functional connectivity strength of specific anatomical regions to the rest of the brain are correlated to the severity of neuroinflammation at month 6.

### Interregional connectivity progressively weakens over disease progression

We first grouped the ROIs into anatomical definitions that mirrored how the brain was dissected for regional neuroinflammation quantification followed by investigating each sub-region’s degree of dysconnectivity to the whole brain. As described in the methods sections, intraregional connectivity was quantified by calculating the mean functional connectivity of all pairwise FC values of regions within an anatomical system/region, and interregional connectivity was quantified by calculating the mean pairwise FC values of regions across an unordered pair of different anatomical systems/regions. Seed to whole brain functional connectivity was quantified by calculating the mean seed ROI’s connectivity to each individual subregion regardless of anatomical system. As observed in **Figure 1A**, there are a greater number of interregional connectivity’s that are significantly different between the 5xFAD and the WT groups relative to the number of intraregional differences. This leads to the notion that specific anatomical regions may be predisposed to disconnecting to the whole brain over disease progression. As observed in Figure 1B, there are multiple regions that exhibit decreased connectivity as a result of disease development at month 6, and the hippocampus is the most featured system to exhibit either interregional or intra-regional connectivity differences between the two groups. As observed in **Figure 1B**, ROIs located within the hippocampal formation and the retro-hippocampus exhibited significantly lower degree of connectivity to the whole brain (p < 0.05, corrected for multiple comparisons). Regions canonically associated with the hippocampus such as the Entorhinal Area, Dentate Gyrus and Subiculum all exhibit significantly lower connectivity to the whole brain and the cortical regions at month 6 (**Figure 1C**). Significance of lower connectivity was quantified using a linear mixed effect model. Similarly, regions featured in the temporal lobe (Temporal association area), parietal lobe (Primary somatosensory area) and hypothalamus (Zona Incerta) also exhibited weaker ROI to whole brain functional connectivity as a result of disease development (**Figure 1D**). The original seed maps and the distribution of subjects for each subregion’s respective connectivity to the whole brain can be found in the supplementary section (**Supplementary Figure 2 - 13**).

**Figure 1:**
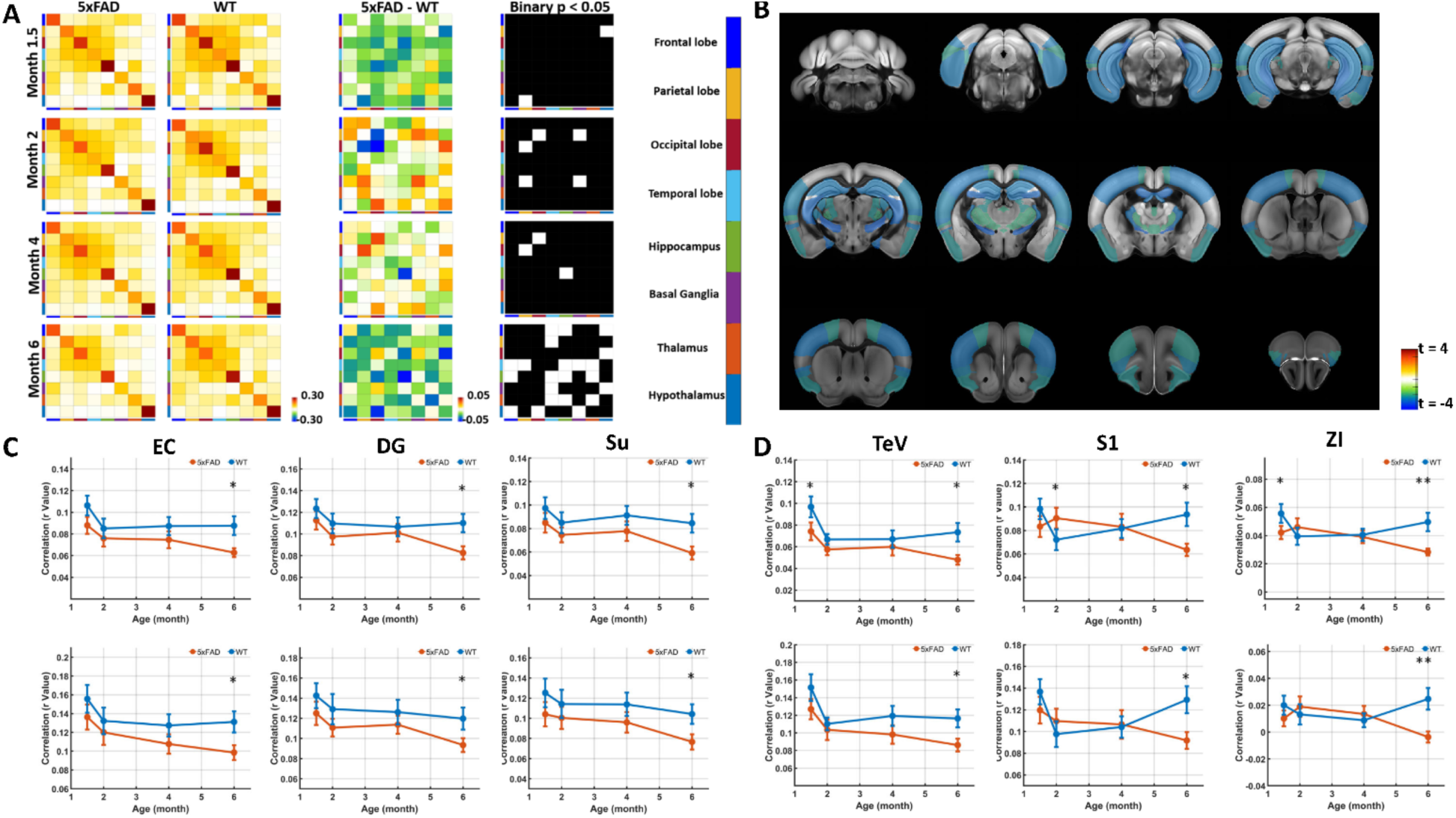
Individual regions progressively disconnect from the brain as disease stage advances. A: Each column denotes a functional connectivity matrix where each cell denotes some metric related to either a region’s mean interregional connectivity with another region, or the mean intraregional functional connectivity, henceforth referred to as the anatomical system connectivity. Each row is a timepoint with month 1.5 being at the top, and month 6 at the bottom. The furthest left column displays the anatomical system connectivity’s for the 5xFAD cohort, the second to the left displays the anatomical system connectivity’s for the WT cohort. The furthest right column displays the binary matrices that denote which interregional/intraregional connectivity is statistically different between the 5xFAD and the WT cohort for the respective timepoint. The second from the right displays the difference matrices, where the group average WT anatomical system connectivity is subtracted from the group average 5xFAD anatomical system connectivity to quantify the direction of difference between the two groups when controlling for timepoint. The color bar for the anatomical systems in the matrices follow the corresponding sequence from the top: Frontal lobe, Parietal lobe, Occipital lobe, Temporal lobe, Hippocampus, Basal Ganglia, Thalamus, Hypothalamus. B: Anatomical mapping of individual regions that feature a statistically significant difference in functional connectivity to the whole brain at month 6. Individual two sample t-tests were run to determine the significance of differences in mean seed ROI to whole brain functional connectivity. Regions shown were controlled for multiple comparisons using false discovery rate. t > 0 values denote that the ROI in the 5xFAD group has a stronger connectivity to the whole the brain, while t <0 denotes that the ROI in the 5xFAD group has a weaker connectivity to the whole brain. C: Select regions from the hippocampus that feature statistically different ROI to whole brain functional connectivity at month 6 (shown in subfigure B) are shown: from left to right is the entorhinal cortex, dentate gyrus, and the subiculum. Top row displays the average ROIs – whole brain connectivity, while the bottom row displays the average ROIs – cortex connectivity. D: Select regions from other regions not in the hippocampus that feature statistically different ROI to whole brain functional connectivity at month 6 (shown in subfigure B) are shown: from left to right is the temporal association area, primary somatosensory area, zona incerta. Top row displays the average ROIs – whole brain connectivity, while the bottom row displays the average ROIs – cortex connectivity.

### Advancing amyloid-β pathology decreases the degree of brain network integrity and modularity

We examined the network’s organizational architecture by quantifying topological measures of assortativity, clustering and global efficiency to assess how disease progression affected the magnitude of network resilience, segregation and integration respectively. First, we observe that there is a global spreading of functional connectivity depression from month 4 to month 6 in the disease cohort at the group level (**Figure 2A**). The group average functional matrices can be found in the supplementary section and demonstrate that the functional differences are not because of negative correlations (**Supplementary Figure 14**). As a result, there is an appreciable increase in the number of sub regional connectivity’s that are lower as a result of advancing amyloid-β pathology, with the genotype effect deemed significant at p < 0.005 for every functional connectivity metric (**Figure 2B**). The individual ROIs that are significant are listed in **Supplementary Table 4.** As mentioned in the methods sections, topological metrics of network theory was quantified using the Brain Connectivity toolbox (Rubinov and Sporns 2010). A linear mixed effect model was used to determine the significance of disease effect at each individual time point. Both 5xFAD and wild-type groups displayed a decline in global brain network organization, reflected by a general trend of reduction in all network topological measures. As implied by the global depression observed in **Figure 2A**, network connectivity is observed to be lower as a result of developing amyloid-β pathology (**Figure 2C**, p_5xFAD:M6-WT:M6_ = 0.035).

**Figure 2:**
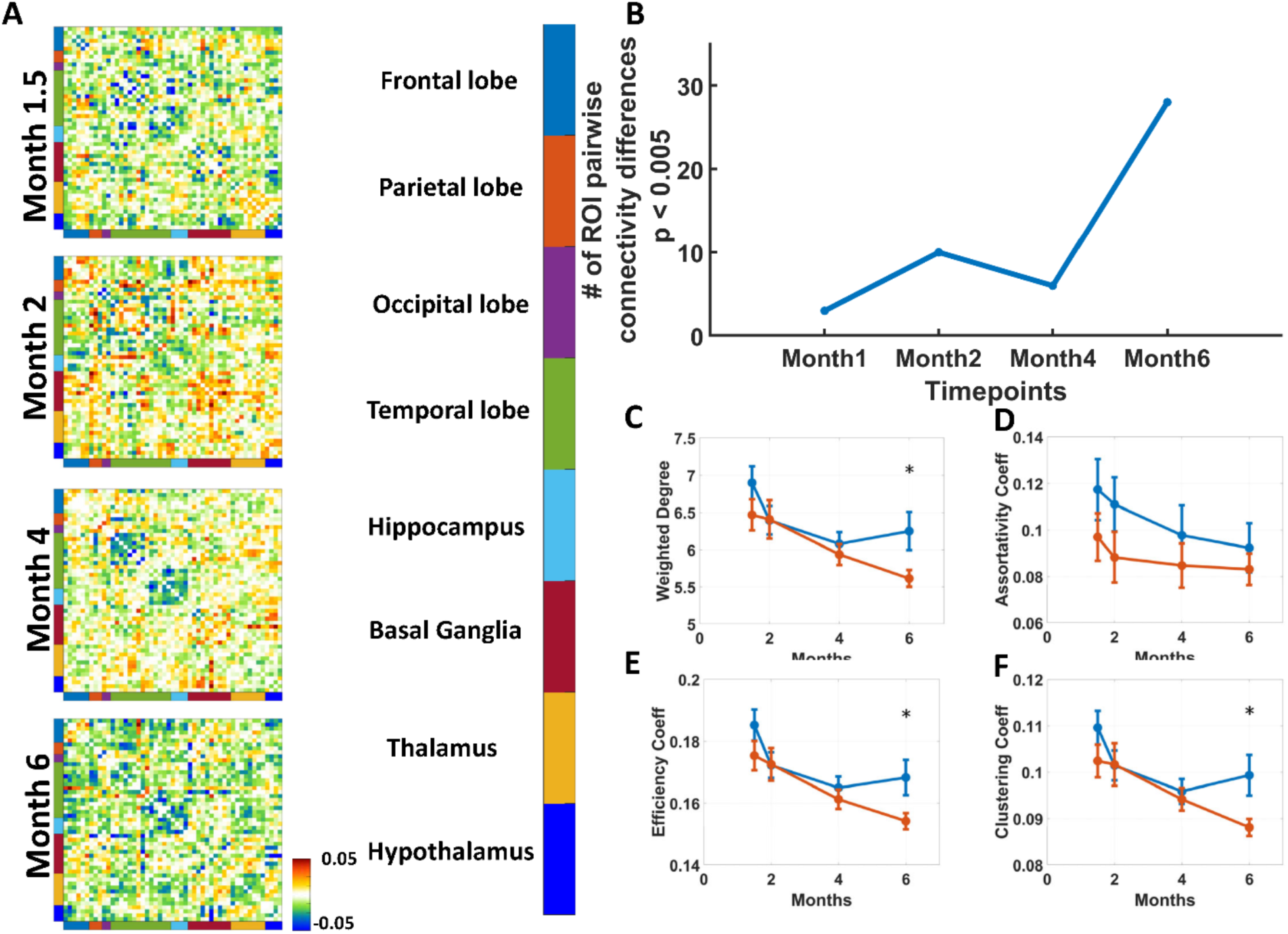
Amyloid-β pathology is associated with weakening metrics of network integrity, modularity and connectivity. A: Difference matrices for each timepoint across 51 ROIs. The ROIs are grouped by anatomical system as denoted by the color bar. The difference matrices are quantified by taking the group average 5xFAD functional matrix and subtracting the group average WT functional matrix from the corresponding 5xFAD matrix at the same timepoint. B: Line graph that qualitatively shows the number of pairwise ROI functional connectivity values that are statistically different. Statistical value is quantified by running a two-sample t test and setting the p value to be 0.005 as the statistical test was not corrected for multiple comparisons. C-F: Line graphs showing the group average graph theoretical metric of interest for both 5xFAD and WT groups with 5xFAD being orange, and WT being blue colored. All statistical tests are two-sample tests and are done to compare month 6 WT versus month 6 5xFAD graph theory metrics. Error bars are standard error of means. The line graphs are as follows C: Global Strength, D: Assortativity, E: Global Clustering, F: Global Efficiency.

Assortativty (network resilience) on the other hand, is observed to have no significant differences for both diseased and healthy aging (**Figure 2D** p_5xFAD:M6-WT:M6_ = 0.569). However, brain networks exhibited a decline in other metrics of network organization such as clustering (network modularity/segregation) and global efficiency (network integration). The decline in both network metrics is appreciably faster in 5xFAD mice, with most topological measures significantly differing between 5xFAD and wild-type mice at Month 6 (uncorrected two-sample t-tests, clustering coefficient: p_5xFAD:M6-WT:M6_ = 0.033; global efficiency: p_5xFAD:M6-WT:M6_ = 0.036, **Figure 2E**, **Figure 2F**). In addition, the later stages of disease development resulted in a lower clustering coefficient compared to early states of disease progression (p_5xFAD:M6-5xFAD:M1.5_ = 0.007, p_5xFAD:M6-5xFAD:M2_ = 0.013) suggesting that amyloid-β neuropathology is driving the underlying biological mechanisms that resulted in depressed metrics of network modularity. The implications of decreased topological metrics of network modularity and integration will be discussed in the discussion section.

### Cytokine signatures are unique to individual brain regions and disease state

Eight brain regions were dissected and assayed to quantify their respective neuroinflammatory signature. Details regarding the dissection and multiplex procedure is listed in the methods. A genotype-specific orthogonalized projection to latent structures regression (OPLSR) model was constructed for every brain region, with the timepoints (month 1.5, month 2, month 4 and month 6) as the response variable and the cytokine concentrations as the covariates. As the models are genotype-specific, any comparison between the two models involving the same brain region can only be done at a qualitative level. Based on our observations from the rsfMRI analysis, we proceeded to investigate 4 of the brain regions that featured disconnecting ROIs from the whole brain as a result of disease progression (**Figure 1B**).

As observed in both **Figure 3A, 3B** (5xFAD column), the cytokine signature for the 5xFAD group for the Temporal and Parietal lobe exhibited a separation of ages, with the older time points (i.e. advanced disease stage) being clustered together. When factoring in the cytokines that make up the inflammatory signature as denoted by their VIP score, we observe that there are less cytokine species driving the diseased aging inflammatory signatures relative to the healthy aging model. Cytokines such as MCP-1 and MIP-1β, MIP-1α, and IL-17A are common cytokines amongst the diseased aging models for cortical regions (**Figure 3E, 3F**, 5xFAD column). The neuroinflammatory signature of diseased aging in the Temporal lobe also featured decreases in IL-12p70, with greater levels of expression in the earlier stages of disease progression (**Figure 3F** 5xFAD column). In contrast to the diseased models, the healthy aging neuroinflammatory signature for the cortical regions is more comprehensive, with each region having more than 9 cytokines making up the signatures (**Figure 3E, 3F** WT column). Additionally, the first latent variable of the healthy aging models captures a higher degree of variance in the predictor space (**Figure 3A, 3B** WT column) relative to the disease models. The significance of this will be discussed in the discussion section.

**Figure 3:**
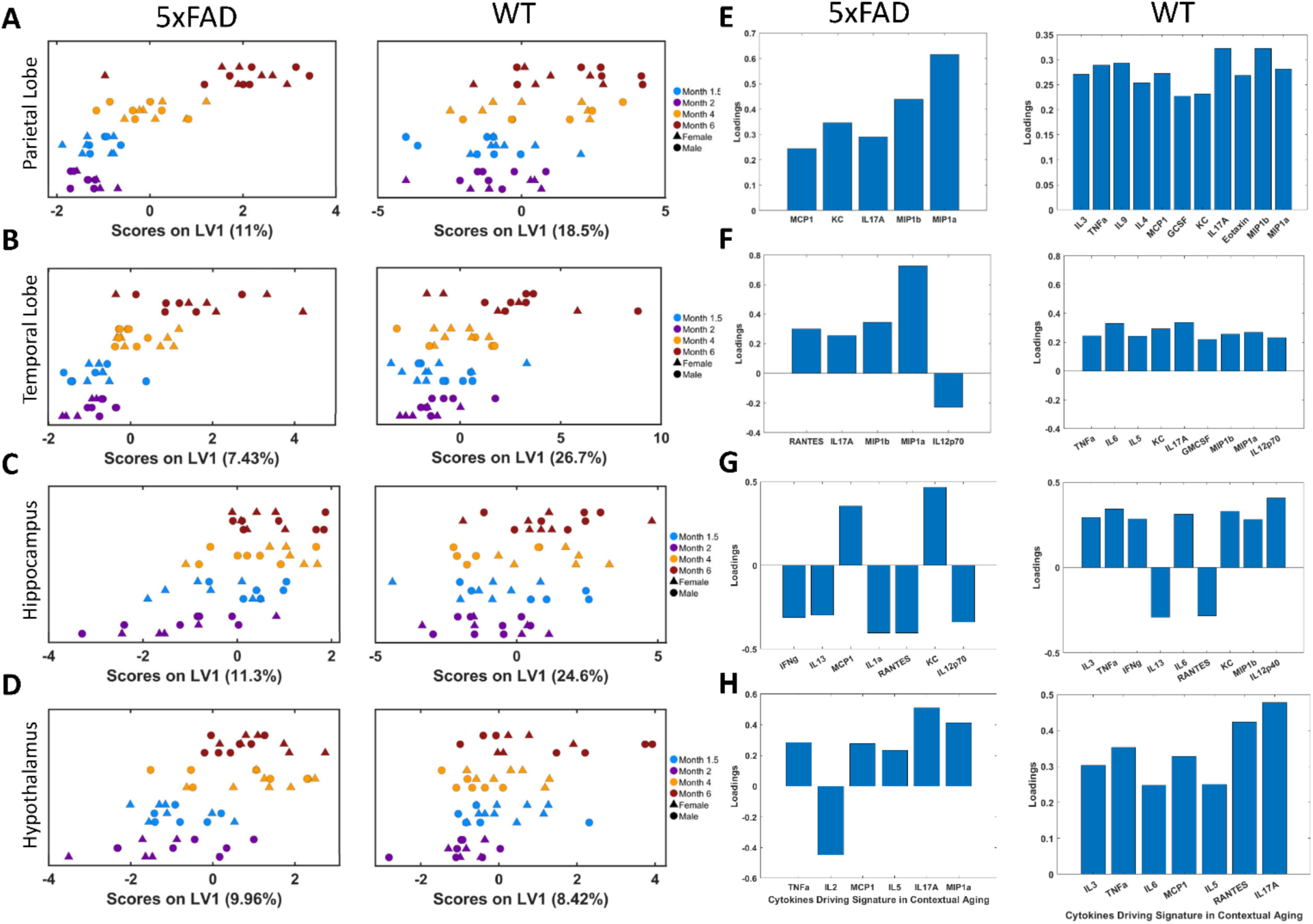
Cytokine signature over disease or healthy aging for select brain regions. A-D: Subject scores relative to the cytokine signature were constructed using the first latent variable of a OPLSR model for the respective brain region and genotype. Each row describes a specific brain region, and each column describes the genotype with the far left being the 5xFAD genotype, second to the left is the WT genotype. The regions are as follows: A: Parietal lobe, B: Temporal lobe, C: Hippocampus, D: Hypothalamus. Points in score plots are color coded by age, with blue describing month 1.5, purple month 2, orange month 4, and red month 6. Sexes are coded with circular points encoding male subjects, and triangular points encoding female subjects. E-H: Loadings that have a VIP score greater than 1 for the corresponding score plots in A-D, with the far right column describing the loadings for the WT model for the respective brain region, and the second to the right column describing the 5xFAD loadings for the respective brain region. The loadings describe the following brain regions in order: E: Parietal lobe, F: Temporal lobe, G: Hippocampus, H: Hypothalamus.

Subcortical regions also displayed a similar feature regarding how the diseased aging model and healthy aging model differed when predicting age. As observed in Figure 3C, 3D, there is a separation of timepoints in the diseased models across the hippocampus and hypothalamus, each one using less than 12% of the variance in covariate space. Unlike the cortical regions, the neuroinflammatory signature of subcortical regions in diseased aging contexts do not share a similar set of cytokines. However, a common feature is that there is an upregulation of neuroprotective cytokines such as IL-2 or IL-12p70 in the earlier stages of disease progression within the subcortical regions (**Figure 3G, 3H**, 5xFAD column). In contrast, the neuroinflammatory signatures in the healthy aging models all featured a greater degree of predictor variance, excluding the hypothalamus (**Figure 3C, 3D** WT Column), and the neuroinflammatory signatures also feature a greater number of cytokine species such as TNF-α, IL-6, IL-5, and RANTES (**Figure 3G, 3H** WT column). The interpretations of these different cytokine signatures will be explored in the discussion section.

### Cytokine signature positively correlates with the magnitude of interregional connectivity for select brain regions

Our prior results demonstrated how both the intersystem connectivity and individual sub regions are disconnecting from the whole brain over disease progression, and that the most noticeable difference is observed at month 6 (**Figure 1A**, **Figure 1B**). Thus, we proceeded to investigate how neuroinflammation is related to a region’s functional connectivity metric. This was achieved in 3 steps: 1) we constructed a projection to latent structure discriminant analysis (PLSDA) model to understand how regional inflammatory signatures differed between disease and healthy states at month 6; 2) anatomical level ROI interregional connectivity (detailed in methods) were quantified; 3) investigated the linear relationship between the functional metrics of interest and the regional neuroinflammation. As our prior results established a pattern of growing dysconnectivity, we tested these models with the hypothesis that a greater disease cytokine signature is negatively correlated with functional connectivity strength (the converse being a stronger healthy cytokine signature is positively correlated with functional connectivity strength).

There are four regions that exhibit a positive correlation between a region’s neuroinflammatory response associated with amyloid-β pathology and a decrease in the respective region’s average intersystem connectivity (**Figure 4**), with those four regions being the Parietal lobe, Temporal lobe, Hippocampus and Hypothalamus. The cytokine signatures for these four brain regions are distinct in nature, but there are some commonalities between them as well. The disease cytokine signature for the two cortical regions both feature the upregulation of MIP-1α and MIP1-β for the 5xFAD group (**Supplementary Figure 17A, 17B**). On the other hand, the hippocampus and the hypothalamus both exhibit their own unique cytokine signature that features increased expressions of cytokines such as IL-1α and KC for the hippocampus (**Supplementary Figure 17C**), and IL-13 and Eotaxin for the hypothalamus (**Supplementary Figure 17D**).

**Figure 4:**
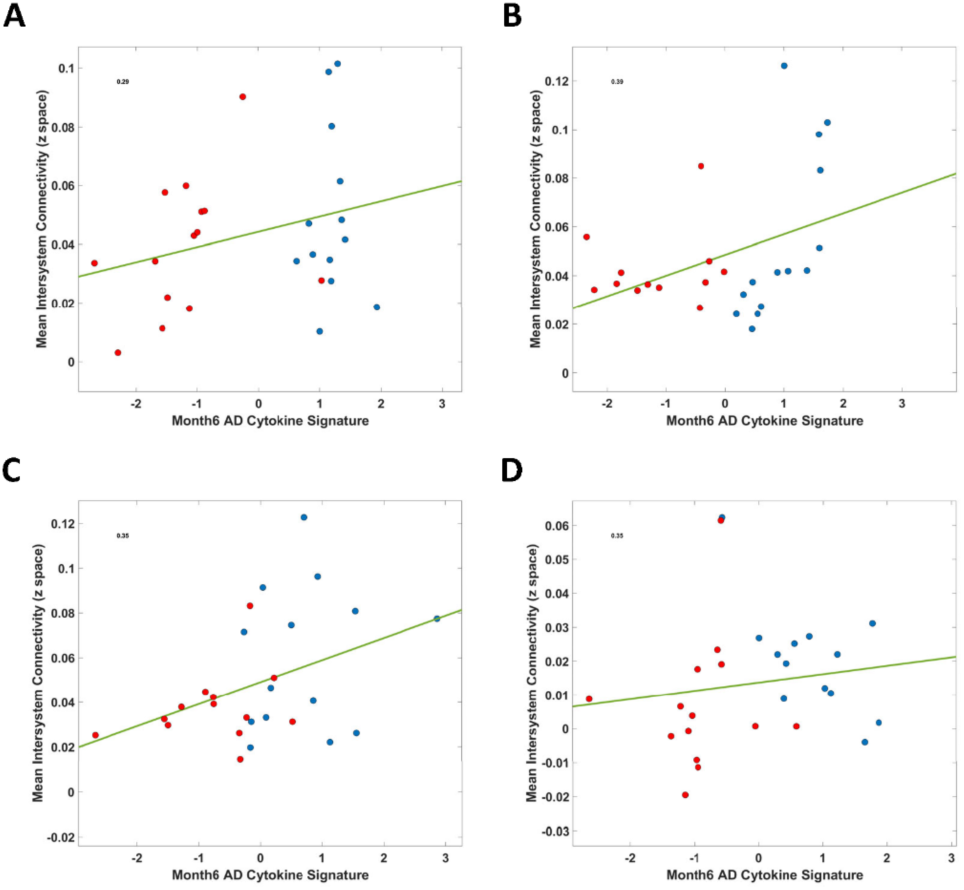
Intersystem functional connectivity negatively correlates with magnitude of disease cytokine signature. Figure 4A-D: Spearman’s correlation with one-tailed t test with the alternative hypothesis stating that the magnitude of intersystem connectivity is negatively correlated with the magnitude of AD cytokine signature at month 6 animals. The spearman correlation results are graphed in the following order: A) Parietal lobe, B) Temporal lobe, C) Hippocampus, D) Hypothalamus. Subjects are color coded to represent AD or WT genotype, with red being AD and blue being WT.

The interrogation of the alternative hypothesis shows that all four brain regions exhibit a positive correlation between a stronger disease cytokine signature and a weaker intersystem connectivity. The Parietal lobe demonstrates the weakest relationship out of the 4 brain regions (r = 0.29, p = 0.067, **Figure 4A**), while the Temporal lobe demonstrates the strongest relationship between disease cytokine signature and interregional functional strength (r = 0.39, p = 0.026, **Figure 4B**). The Hippocampus and Hypothalamus both demonstrate a robust relationship between the two metrics as well, with both the Hippocampal and Hypothalamic model demonstrating a relatively strong relationship between cytokine signature and interregional connectivity strength at r = 0.35 (**Figure 4C**, **Figure 4D**).

## Discussion

We investigated how regional neuroinflammation and functional connectivity properties of the brain are related during the development of Alzheimer’s disease pathology. To accomplish the aforementioned objective, we quantified resting state functional connectivity (RSFC) and the regional immune-milieu to determine the relationship between the two modalities. Awake resting state fMRI was utilized to quantify RSFC as it enabled the measurement of basal regional neuronal activity without the confounding variable of anesthesia. Functional connectivity metrics were quantified by correlating the BOLD time courses of different ROIs, with the seed ROI to whole brain metric being the mean connectivity of the seed ROI to every other ROI in the brain. Inter and intra system connectivity metrics were measured by grouping ROIs into anatomical sets followed by quantifying the mean connectivity between sets or within sets respectively. Network analysis of the functional connectivity matrices was achieved by calculating graph theory metrics by treating the matrices as adjacency matrices and quantifying topological features related to the matrices. Multiplexing of cytokine species was utilized to quantify regional cytokine signatures by multiplexing individual brain regions that were dissected from one hemisphere of the brain. Multivariate analysis was conducted on the cytokine dataset to infer the cytokine signature for healthy and diseased aging contexts. Lastly, multivariate analysis on advanced disease stage subjects yielded the AD cytokine signature that delineated how the immune-milieu is different between healthy and disease animals. The subsequent AD cytokine signature is then correlated with functional connectivity metrics, with four brain regions exhibiting a relationship where a stronger disease cytokine signature is correlated with a weaker degree of inter-system connectivity.

### Subregions in the Parietal lobe, Temporal lobe, Hippocampus and Hypothalamus exhibit weaker connectivity to the whole brain as AD develops

The grouping of individual ROIs into their respective anatomical systems and the subsequent analysis into intra and inter-regional functional connectivity revealed that disease progression is correlated with weaker intra and inter-regional connectivity (**Figure 1A**). We observe that the overall strength of anatomical interregional and intra-regional connectivity strengths is positive (**Figure 1A**, 5xFAD and WT column), and that the differences between the two groups are driven primarily by a weaker degree of functional connectivity, instead of the difference being driven by one group featuring negative correlations (Figure 1A, 5xFAD-WT column). Thus, with the prior two observations in mind, we saw that month 6 featured the greatest number of differences regarding interregional and intraregional connectivity’s. The hippocampus is the most prominent anatomical system to be featured in functional connectivity metrics, with diseased subjects expressing lower connectivity between ROIs in the hippocampus and the temporal lobe, parietal lobe, and hypothalamus at month 6 (Figure 1A, Binary column). Alongside the findings of regional connectivity of anatomical systems, our results also illustrate how individual brain regions in the hippocampus and cortical areas exhibit lower connectivity to the whole brain at the most symptomatic time point (**Figure 1B, 1C**). The manner in how weaker functional connectivity values first are observed within the hippocampal network at month 4 (**Figure 1A**, Binary Column) before observing multiple regions exhibiting weaker connectivity at month 6 mirrors the trajectory of amyloid-β pathology, as the hippocampus and retro hippocampus are the first areas to be affected in the pathological development timeline, with the cortical areas affected next (Thal et al 2002, Braak et al 2011, DeTure and Dickson 2019). Some of the hippocampal regions implicated in our study as a result of progressive amyloid-β pathology have been documented to play a role in memory forming and retrieving processes such as Field CA3, Dentate Gyrus (**Figure 1C**) (Chadwick et al 2014, Hainmueller and Bartos 2020), and the Temporal association area located in the temporal lobe (Villa and Fuster 1992, Sakai and Miyashita 1993).

While the hippocampus is heavily implicated in our results (and in existing literature), there were brain regions in other anatomical systems that also exhibited weaker connectivity as a result of disease proteinopathy. The other two cortical regions that exhibited weaker connectivity to the whole brain in the 5xFAD cohort at month 6 are the gustatory area and Primary somatosensory area (**Figure 1D**), regions defined to be in the Insular cortex (and as part of the Temporal lobe for anatomical system analysis) and Parietal lobe respectively. The disconnecting of the gustatory area over disease progression in this study reflects some of the recent findings regarding how gustatory dysfunction is correlated with disease pathology (Suto et al 2014, Martin et al 2018), as amyloid-β plaques can build up in the insular cortex during the early stages of disease development (Bonthius et al 2005). A surprising finding from the study is how the Primary somatosensory area’s functional connectivity is implicated as a result of disease progression. This is because one of the results from Palmqvist et al 2017 was that the sensorimotor cortex is a region that does not accumulate a significant amount of amyloid-β in the preclinical to middle stages of AD. However, there have been other studies that have documented how cognitive functions related to the sensorimotor network are affected in AD (Wiesman et al 2021, Martínez-Pernía et al 2023), and that the 5xFAD mouse model exhibits an accumulation of amyloid-β in the sensorimotor network (Plachez et al 2023). The interpretation that could be made is that regions that experience amyloid-β pathology such as the hippocampus, the medial temporal lobe (Thal et al 2002, Braak et al 2011, DeTure and Dickson 2019), insular cortex (Bonthius et al 2005), and the parietal lobe (Plachez et al 2023) is linked to the weakening of functional connectivity to the brain. This helps explain why certain cognitive processes are impaired over the course of progressive amyloid-β pathology as specific brain regions are selectively more impacted from disease pathology.

Furthermore, the Zona Incerta was also disconnecting from the whole brain at the most symptomatic disease state (**Figure 1D**). The Zona Incerta has been hypothesized to have some global function (Mitofanis 2005), with the region being featured in a diverse set of different cognitive functions such as locomotion (Perier et al 2002) and attention (Tait et al 2017). The implications of a weakening connectivity for the Zona Incerta can result in changes to behavior, where the suppression of the rostral part of the Zona Incerta lead to increases in freezing time in a rodent model undergoing fear conditioning experiments (Chou et al 2018). However, the reason as to what is driving the dysconnectivity of the Zona Incerta can not be strongly attributed to amyloid pathology, as it was found that there were low levels of amyloid plaques in the thalamus and marginal levels in the hypothalamus in 9-month 5xFAD mice (López-Gambero et al 2021, Tsui et al 2022). Therefore, it is possible that the dysconnectivity of the region is driven by its connections to regions in the brain that are directly affected by disease pathology. It has been shown that the subthalamic (defined as hypothalamic in the paper) subregion has projections to disease affected subregions in the hippocampus (Zhang et al 2024) and to the primary somatosensory cortex (Nicolelis and Lin 1995). Thus, the Zona Incerta is an example of how a region can be indirectly affected by disease pathology, with the consequence being becoming functionally isolated and unable to execute basal functions that underlie a normal system.

### Developing amyloid pathology disrupts brain network characteristics

There are widespread differences between the 5xFAD and WT groups over aging, with the 5xFAD group exhibiting weaker functional properties relative to the WT group as disease pathology develops (**Figure 2A**). This is confirmed by **Figure 2B**, where we observe an inflection point regarding the number of differing functional connectivity’s that are statistically significant between month 4 and month 6. The increasing number of ROIs that are different between the two time points, and how widespread those differences are suggest that there is a global depression of network strength in the diseased brain. Thus, we proceeded to quantify 4 different network metrics that help us understand the network properties of the diseased brain. The 4 metrics of interest are: global strength, assortativity, global clustering, and global efficiency. All of the global metrics are shown to be decreasing in magnitude over the aging for both 5xFAD and WT groups, with the 5xFAD group displaying a greater decrease in the global metrics at month 6 (**Figure 2C, E, F**). The decrease in a network’s global strength suggests that there are multiple nodes within the network that are becoming less functionally connected, resulting in a decrease in a network’s integrity. Contrasting the difference in connectivity, the similar profiles for assortativity in the 5xFAD and WT groups suggest that the two networks do not lose resilience relative to each other as a result of disease progression (**Figure 2D**). The implication is that regardless of how the brain network ages, nodes that have specific properties will tend to connect to other nodes of similar properties in a manner that is indifferentiable between 5xFAD and WT groups.

The corresponding decrease in the global clustering coefficient suggests that the network’s ability to segregate itself into modules is impaired as a result aging, and that the buildup of amyloid pathology accelerates the loss of network segregation/modularity (**Figure 2E**). Modularity is an important network metric to quantify as it signifies how well a network can define groups of nodes, with the perception that a higher degree of modularity corresponds to specialized functional networks for specific cognitive processes (Bavassi et al 2019). Thus, the implication of a lower degree of modularity is the decreased ability to execute cognitive tasks (Wang et al 2021, Zuberer et al 2021, Cohen and D’Esposito 2016). Similar results have been observed in human studies as well, where advanced disease pathology is associated with lower levels of functional segregation (Ng et al 2021, Cai et al 2020).

The same pattern was also observed for global efficiency, with global efficiency decreasing in both disease and healthy aging, and that the fully symptomatic timepoint resulted in a significant difference between the two groups (**Figure 2F**). Global efficiency is a metric of network integration, and it has been reported that expanded cognitive ability coincides with higher levels of network integration as it measures how well information can travel from one node in the network to every other node in the network (van den Heuvel et al 2009). Therefore, the inability to integrate information throughout the network in the 5xFAD group relative to the WT group suggests that it is the developing amyloid pathology that drives the loss of network topological features. Consequently, cognitive deficits arise due to an inability to segregate into modules and the inability to integrate functional information from disparate brain regions.

Our results do show that there is a decrease in network metrics from month 6 relative to the earlier timepoints. Distribution of subjects across all graph metrics show that on average, there are differences between the 5xFAd and WT groups when not factoring in sex (**Supplementary Figure 15**). It has been reported that aging is correlated with decreased cognitive functionality, and levels of network integration and segregation (Crowell et al 2020, Varangis et al 2019). As such, the decrease of network metrics in our WT group is to be expected, and that the lower values in network metrics in our 5xFAD group strongly suggests that it is the presence and buildup of amyloid-β that led to the dysfunctionality of the brain network in the diseased aging group. The significance of this interpretation is that as Alzheimer’s disease progresses, the brain becomes less capable of executing complex cognitive tasks (Cohen and D’Esposito 2016, Zuberer et al 2021), whilst the healthy aging brain is able to recover, giving one potential reason why healthy elderlies do not experience the same degree of cognitive impairment to Alzheimer’s patients. Evidence suggesting the aforementioned claim can be found by examining healthy aging literature. It has been reported that healthy aging brains may be able to compensate for age-related functionality declines (Ji et al 2018, Larivière et al 2019), and that the diseased brain is unable to undertake such processes given the breadth of tissue damage as a result of chronic inflammation due to disease pathology (Bader et al 2023, Wood et al 2015, Braak et al 2006, Wang et al 2004, Jack et al 1999, Jayaraman et al 2021, Kwon and Koh 2020).

### Healthy aging cytokine signatures are more comprehensive and neuroprotective relative to diseased aging signatures

We dissected and assayed a number of different brain regions from one hemisphere of the brain for both 5xFAD and WT groups to understand what the neuroinflammatory signatures are for different brain regions over aging. Our results show that the cytokine signature for diseased aging is qualitatively different from healthy aging, with the number of cytokines having a VIP score greater than 1 to be lower in the 5xFAD aging cohort relative to the WT cohort across the 4 brain regions (**Figure 3**).

Cytokines with a VIP score greater than 1 are cytokines that are found to have a high degree of covariation with the response variable, which in this case is time. Additionally, the first latent variable in the disease aging models, excluding the hypothalamus, captured less variation in the covariate space relative to the healthy aging models (**Figure 3A-D**). Coupling the two observations suggest that the cytokines featured in a diseased aging neuroinflammatory signature have a high magnitude of covariation with disease state, and are driving the separation across the different disease states. Thus, when we factor in the specific cytokines featured in the disease models, we observe that the cytokines are predominantly inflammatory in nature (**Figure 3E-H**).

Some of cytokines expressed in diseased aging are commonly found in AD contexts such as MIP-1α, and MIP-1β (Passamonti et al 2019, Liu et al 2014). These cytokines are featured in all cortical regions (**Figure 3E-F**, **Supplementary Figure 16**), while MIP-1α and not MIP-1β is featured in the Thalamus, Hypothalamus and Striatum disease models (**Figure 3H**, **Supplementary Figure 16**). Furthermore, other cytokines featured in the diseased aging models confirm that the cytokine signatures for diseased aging are predominantly inflammatory in nature. By examining the cortical regions alone, we observe that cytokines that covary positively with diseased aging are typically associated with inflammation (**Figure 3E, 3F**, **Supplementary Figure 16**), such as RANTES (also known as CCL-5), MCP-1, IL-17A, KC (also known as CXCL-1), and the aforementioned MIP-1α and MIP-1β (Sui et al 2004, Sui et al 2006, Vlkolinský et al 2004, Bhavsar et al 2015, Marciniak et al 2015, Ivanovska et al 2020). A notable exception is how IL-10 covaries positively with diseased aging in the occipital lobe (**Supplementary Figure 16**) as IL-10 has been reported to have neuroprotective functionalities but is also highly co-expressed with canonically known proinflammatory cytokines (Saraiva and O’Garra 2010). One interpretation that could be made is that there is an inflammatory response in the Occipital lobe due to amyloid-β pathology, but the negative consequences are mediated due to an increased expression of IL-10. This is reflected in our PLSR score plots with little separation between groups (**Supplementary Figure 16**), and from our functional results, where we observed no differences in the intra-regional connectivity of the Occipital lobe between the diseased aging and healthy aging cohorts (**Figure 1A**, Binary Column), and that no ROI within the Occipital lobe exhibited a weaker connectivity to the whole brain at the most symptomatic state of the study (**Figure 1B**).

Another interesting observation for the cortical diseased regions is that the frontal lobe and the temporal lobe have cytokines that are negatively correlated with disease state, with IL-2 having a higher expression levels at the earlier stages of disease progression for the frontal lobe (**Supplementary Figure 16**), and IL-12p70 having the same feature for the Temporal lobe (**Figure 3F** 5xFAD column). What makes these features stand out is that IL-2, and IL-12p70 have been reported to have some neuroprotective effects, with IL-2 reported have a regulatory role in inflammation and neuronal activation (Yshii et al 2022, Alves et al 2017, Meola et al 2013), and IL-12p70 reported to regulate the degree of microglia related neurodegeneration (Andreadou et al 2023). Additionally, IL-2 and RANTES have been reported to have a have a synergistic relationship to promote an optimal inflammatory response (Weiss et al 2009, Parisi et al 2020). Thus, it could be inferred that the downregulation of RANTES had some neuroprotective aspect in the region’s cytokine signature (**Supplementary Figure 16**). This in turn may have delayed the negative effects of developing amyloid-β pathology, which is illustrated through our functional connectivity data. The frontal lobe was a region that did not exhibit a large number of ROIs that had weaker connectivity to the whole brain as a result of disease progression (**Figure 1B**). The same can not be said about the Temporal lobe as RANTES was upregulated as disease stage progressed (Figure 3E 5xFAD column), which may indicate how the underlying immune-milieu is driving the observed differences in the number of individual ROIs that are disconnecting from the whole brain at the symptomatic state (**Figure 1B**).

Subcortical regions (Hippocampus, Striatum, Thalamus and Hypothalamus) do not share the same exact neuroinflammatory profile for diseased aging as cortical regions but do seem to have the same consequences. Inflammatory cytokines such as MCP-1, and KC were upregulated for the Hippocampus and RANTES was downregulated **(Figure 3G** 5xFAD column). Inflammatory cytokines such as MIP-1α, IL-17A were upregulated in the other three subcortical regions, alongside with upregulation of specific cytokines such as IFN-γ for Striatum and Thalamus **Supplementary Figure 16**), and MCP-1 for Hypothalamus (**Figure 3H** 5xFAD column). Similar to the Frontal lobe, the Hippocampus, Thalamus and Hypothalamus had a downregulation of IL-2 and RANTES (**Figure 3H** 5xFAD column **Supplementary Figure 16**), which would suggest that there was an initial neuroprotective aspect to the earlier cytokine signatures but that as amyloid pathology persisted, this neuroprotective feature went away and essentially enabled the subsequent neuropathological issues observed in Alzheimer’s disease subjects. The Thalamus and Hypothalamus exhibited similar functional connectivity properties to the Frontal lobe, with only one region in their respective anatomical systems exhibiting weaker connectivity to the whole brain at the terminal symptomatic state (**Figure 1B**). This is in line with the expected neuropathology of Alzheimer’s disease, with amyloid-β buildup in subcortical regions to occur at the later stages of disease development (Thal et al 2002, Braak et al 2011, DeTure and Dickson 2019).

However, this does not explain how the Hippocampus has a similar disease and healthy aging cytokine profile even though it is the first region to be affected (Thal et al 2002). This in fact could be explained by how it is RANTES that is driving the cytokine signature and not IL-2 as RANTES (also known as CCL5) plays a significant role in mitochondrial stability and that the lower expression levels of RANTES may result in neuronal dysfunction (Ajoy et al 2021). Moreover, TNF-α is another proinflammatory cytokine (Heppner et al 2015) that is upregulated in the diseased aging cytokine signature of the hippocampus (**Figure 3G**. 5xFAD column). The coupling of TNF-α alongside other proinflammatory cytokines and the downregulation of RANTES may have resulted in the hippocampus being affected earlier in disease development even though the Hippocampus shares a similar cytokine profile (with regards to RANTES and IL-2 having negative covariation with disease stage) with the Frontal lobe, Thalamus and Hypothalamus. Once again, this is reflected in the functional data where the 5xFAD cohort exhibited greater differences in two different rsFC metrics: intra-regional connectivity at month 4 and 6 (**Figure 1A**, Binary column), and the number of ROIs in the hippocampus that had weaker connectivity to the whole brain at the symptomatic state (**Figure 1B**).

The healthy aging profiles across all regions (both cortical and subcortical) excluding the Hypothalamus are similar in the sense that they all feature substantially more cytokines in the neuroinflammatory profile relative to their respective diseased aging models (**Figure 3E-H**). For the cortical regions the heathy aging models featured proinflammatory cytokines such as TNF-α, MIP-1α, MIP-1β, and CCXL1 (Bhavsar et al 2015, Marciniak et al 2015, Wakabayashi et al 2021), but also featured at least one anti-inflammatory cytokine such IL12-p70 (Andreadou et al 2023) for the Frontal, Occipital, and Temporal cortex (**Figure 3F**, WT column, **Supplementary Figure 16**), and IL-4 (Francos-Quijorna et al 2016, Fenn et al 2014) for the Parietal Cortex (**Figure 3E**, WT column). The Thalamus and Striatum share a similar feature where IL-10 is upregulated in the Striatum and IL-12p70 is upregulated for Thalamus (**Supplementary Figure 16**). The observation that these models capture more variation in the covariate space suggests that there isn’t a specific subset of cytokines that are driving the differences in cytokine profiles across healthy aging groups. Consequently, the interpretation that could be made is that healthy aging does lead to some inflammatory response, but not to the same degree as the 5xFAD cohort. This interpretation is in line with other studies that have examined how aging and inflammation are related, with it being reported that microglia in the aged brain have higher expression levels of proinflammatory cytokines (Norden and Godbout 2013). Additionally, inflammation itself is not necessarily detrimental as the co-expression of neuroprotective cytokines such as IL12-p70, IL- and IL-4 suggest that the inflammatory profile may not be neuroinflammatory in nature and may be more so neuroprotective or neuromodulating instead.

Therefore, when we couple the healthy aging inflammatory results with our quantification of global clustering and global efficiency, we can draw a future hypothesis to be tested. Our data suggests that the healthy aging immune-milieu enables a neuronal environment that allows for neuronal compensation at a system level, which allows for brain networks in healthy aging subjects to recover network modularity and efficiency.

### Magnitude of diseased cytokine signature is negatively correlated with the strength of intersystem connectivity

We have observed that diseased aging in the assayed regions has a more proinflammatory cytokine signature relative to the healthy aging models (**Figure 3**), and that ROIs are exhibiting weaker connectivity to the whole brain at the symptomatic disease state which is also known as month 6 for the 5xFAD cohort (**Figure 1**, **Figure 2**). Given these prior observations, we proceeded to quantify how the cytokine signature differed between 5xFAD and WT cohorts at month 6 in order to investigate whether specific cytokine signatures are correlated with weakening functional connectivity as a result of advancing amyloid-β pathology. As such, we have observed 4 brain regions (Parietal lobe, Temporal lobe, Hippocampus and Hypothalamus) where the difference in cytokine signatures between month 6 5xFAD and WT animals is correlated with the respective brain region’s weakening intersystem functional connectivity to the whole brain (**Figure 4**).

The cytokine signature that differentiates between the Parietal lobe in the 5xFAD cohort and the healthy aging cohort is driven by 4 cytokines that have higher levels of expression levels for the 5xFAD group. These cytokines are KC, MIP-1α, MIP-1β, and IL-12p40 (**Supplementary Figure 17**). KC, MIP-1α, and MIP-1β are chemo-attractants for immune cells, which suggests that the 5xFAD group is experiencing a heightened inflammatory response in the Parietal lobe relative to the WT group. The same feature can be applied to the Temporal lobe, which has a heightened expression levels of IL-2, MIP-1α, MIP-1β for the 5xFAD group, and an upregulation of IL-12p70 and IL-5 for the WT group (**Supplementary Figure 17**). The correlation between two regional neuroinflammatory profiles and their relevant intersystem functional connectivity would suggest that developing amyloid-β pathology weakens intersystem connectivity (**Figure 4A** pVal = 0.067, **4B** pVal = 0.026). The observation that the Parietal lobe and Temporal lobe have both disconnecting functional connectivity’s and an inflammatory response mirror the expected disease pathology. The Temporal lobe is one of the cortical areas heavily affected by Alzheimer’s disease pathology (Thal et al 2000, Thal et al 2002, Braak et al 2011), and the 5xFAD model exhibits a buildup of amyloid-plaques in the Parietal lobe (Plachez et al 2023). This result provides evidence that neuroinflammation can disrupt a set of neuron’s abilities to coordinate activity, which can lead to an alerted cognitive performance (Yamamoto et al 2019, Nguyen et al 2020, Akiyohsi et al 2018).

The Hippocampus is an area that was expected to exhibit a correlative relationship between the region’s functional connectivity and neuroinflammatory profile due to how early the region is affected in disease pathology. Whilst we do observe the correlation between the two modalities, with a greater inflammatory profile negatively correlated with the strength of connectivity between the Hippocampus and every other anatomical system (**Figure 4C**, pVal = 0.039), the cytokine signature illustrating the difference in a regional’s inflammatory response consisted of species that were not expected; mainly IL-2 and RANTES being upregulated for the 5xFAD group (**Supplementary Figure 17**). As mentioned before, IL-2 and RANTES have a synergistic relationship in promoting an optimal inflammatory response (Weiss et al 2009, Parisi et al 2020), and so observing the two being upregulated in the 5xFAD group would suggest that the degree of glial activation is being modulated. However, we also observe that IL-1α and KC are upregulated and IL-12p70 is downregulated for the 5xFAD group (**Supplementary Figure 17**), suggesting that the cytokine signature suggesting that the differences between the diseased and healthy aging groups is primarily driven by the degree of inflammation, with the 5xFAD group having a greater degree of neuroinflammation than the WT group in the Hippocampus.

The hypothalamus was a surprising result as the 5xFAD model does not have a noticeable buildup of amyloid-β in the Hypothalamus at month 6 (López-Gambero et al 2021, Tsui et al 2022). However, our results show that the hypothalamus as a whole is disconnecting from the whole brain as a result of 5xFAD pathology, and that there is a corresponding inflammatory profile that correlates with the weakening intersystem connectivity (**Figure 4D**, pVal = 0.038). Similar to the other three regions, the cytokine signature of the Hypothalamus that separated the 5xFAD and the WT subjects was inflammatory in nature, with IL-13, Eotaxin and MIP-1α being upregulated for the 5xFAD group, and RANTES and KC being upregulated for the WT group (**Supplementary Figure 17**). One potential reason as to why the functional connectivity of the region was affected by inflammation is that the Zona Incerta was observed to have a weaker connectivity to the whole brain at month 6 (**Figure 1D**) and that the Zona Incerta has projections with regions in the dorsal hippocampus such as the Dentate gyrus and Field CA 1 (Zhang et al 2024). It is likely that the Hypothalamic ROI’s had weaker connectivity due to its neuronal projections to regions in the Hippocampus.

In summary, each region had their own unique cytokine signature that explained how the neuroinflammatory environment between the 5xFAD and WT animals differed for the 4 aforementioned regions. The cortical signatures are similar as they both feature MIP-1α and MIP-1β as one the major cytokines that are different between the 5xFAD and WT groups, but the subcortical regions have a more distinct profile, with the Hippocampus not featuring any MIP-1 family chemokines and the Hypothalamus featuring two cytokines not featured in the other three regions (IL-13 and Eotaxin). Whilst we speculate that all of the cytokine signatures point towards an inflamed environment, which can explain the weakening functional connectivity due to the presence of activated microglia (Yamamoto et al 2019, Nguyen et al 2020, Akiyohsi et al 2018), it must be stated that the nature of each environment may be different due to the different cytokine species being elevated in their respective regions.

### Limitations

There are limitations to our findings and the dataset presented here. Ultimately, we have used an animal model of Alzheimer’s Disease. Thus, our dataset was formed using a highly aggressive mouse model that best replicates familial AD, and not sporadic AD. In more ways than one, any interpretation from the collected data can only be done after considering that it was an abnormal accumulation of human mutant amyloid-β that drove disease pathology. It is likely that sporadic AD may have a different timeline in terms of AD neuropathology and proteinopathy. Moreover, no animal behavior was quantified with the animals in this dataset. Animal behavior would have provided a more comprehensive insight as to how changes to neuronal behavior due to neuroinflammation can affect both resting state functional connectivity and cognitive performance. Additionally, the spatial resolutions of the of the cytokine data and the fMRI data do not match up, with individual ROIs in the fMRI analysis being substantially smaller than any of the brain regions assayed. This leads to the inability to quantify how individual regions connectivity is influenced by an inflammatory profile, and whether a specific set of regions are driving the overall functional dysconnectivity. Lastly, the mice that had their brains dissected could only have one timepoint. It is not currently possible to create a true longitudinal cytokine dataset that mirrored the experimental time points of our imaging data. If possible, a longitudinal cytokine dataset alongside a longitudinal imaging dataset would control for subject variances, a notable concern when working with animal models.

### Summary

In this paper, we have quantified regional neuroinflammatory profiles that correlate with the respective strength of the region’s functional connectivity to the whole brain. We report that the entire brain’s network loses topographical features as amyloid-β pathology advances, and that individual regions within the Hippocampus, Temporal lobe, Parietal lobe and Hypothalamus are disconnecting from the whole brain over disease progression. Cytokine signatures of these regions were quantified using multivariate models that extracted key cytokine and chemokine species that delineate diseased aging or healthy aging, and that these corresponding models elucidated how each region was progressively getting inflamed and that the diseased aging group was predominantly neuroinflammatory, whilst the healthy aging models showed a more measured inflammatory environment. In addition to the aforementioned models, we also ran a multivariate model to understand how the neuroinflammatory environment differed at month 6 and what their relationship is with the strength of functional connectivity. What we report is that the Hippocampus, Parietal lobe, Temporal lobe and Hypothalamus exhibit a correlative relationship between neuroinflammation and the corresponding strength of functional connectivity, with diseased animals having a weaker connectivity alongside a more inflamed environment. The two cortical regions and the Hippocampus region fall-in-line with existing literature as amyloid-β pathology does affect these regions at month 6. However, the Hypothalamus was a surprising result as it had been reported that there is no significant buildup of amyloid-β in the region at month 6, but it still featured a more inflamed environment in the 5xFAD cohort and a weaker connectivity. What this suggests is that there are unique cytokine signatures that underlie the strength of a region’s functional connectivity, and that these 4 regions are vulnerable to developing disease pathology. Therefore, these regions are potential targets for future therapeutic strategies that aim to recover a region’s functionality by mediating the neuroinflammatory environment with the objective to restore the strength of functional connectivity to the whole brain.

## Acknowledgements

This work was supported by R21AG068532 from the National Institute on Aging (EAP/NZ), grant TSF41000095613 from the Pennsylvania Department of Health using Tobacco CURE Funds (EAP/NZ), and start-up funds from Penn State College of Medicine Departments of Neurosurgery and Pharmacology (EAP). DCC and MKK were supported by training fellowship T32NS115667 from the National Neurological Disorders and Stroke. All funding agencies disclaim responsibility for any analyses, interpretations, or conclusions.

The authors would like to acknowledge the Huck Institutes’ High-Field Magnetic Resonance Imaging Core Facility (RRID:SCR_024461) for use of the Bruker Biospec 70/30.

## Contributions

Dennis Chan, Nanyin Zhang, Elizabeth Proctor designed the experiments, with contribution from Kevin Turner. Dennis Chan conducted the surgical implantation, awake animal imaging, brain dissections, and the primary analysis. Samuel Cramer conducted secondary analysis. Chaemin Kim, Rachel Kang, and Madison Kuhn conducted the Luminex experiments and genotyping. Lynne Beidler conducted genotyping and brain dissection. Dennis Chan designed the primary equipment. Said Hayreddin Ünsal and Samuel Cramer designed the secondary equipment. Thomas Neuberger designed the imaging protocol. Dennis Chan, Nanyin Zhang, and Elizabeth Proctor edited the paper. Nanyin Zhang and Elizabeth Proctor conceptualized the work and acquired the funding.

## Notes

### Competing Interest Statement

The authors have declared no competing interest.

### Summary of Updates

additional analysis, revised figures, improved text.

