## Supplementary Materials for "Cytokine expression patterns predict suppression of vulnerable neural circuits in a mouse model of Alzheimer’s disease"

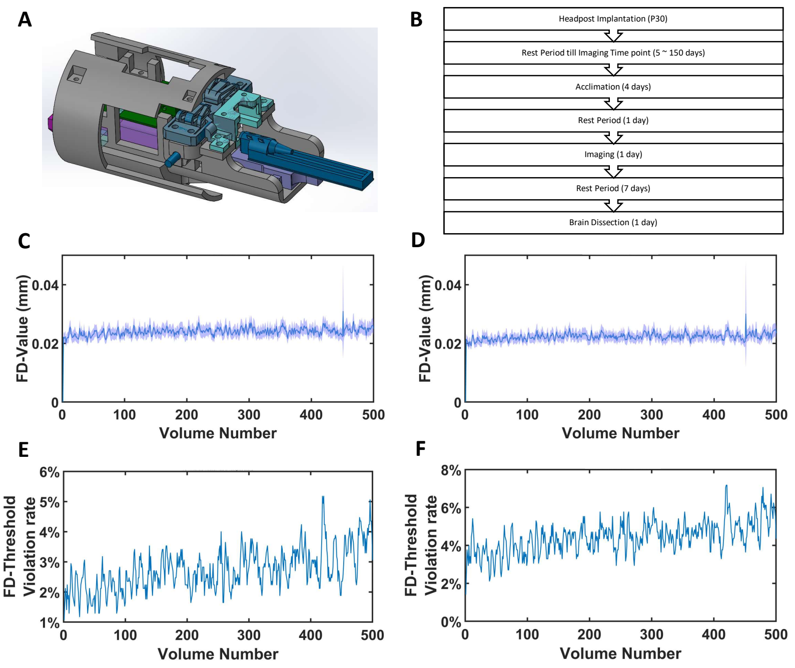

Supplementary Figure 1: A) 3D rendition of the custom mouse restrainer that was designed and used for awake resting state fMRI. Colored pieces that are not gray designate which parts are modifiable to accommodate for different sized mice. The dark blue highlighted nose cone has angled perforations on the dorsal side of the piece to allow gaseous diffusion without significant loss of changes in air pressure during breathing such that the respiration rate can be quantified without having the mice inhale recycled air. The dark blue highlighted ear bars help keep the head steady during imaging. The light purple highlighted piece is the nosecone mount where different heights allow for proper fitting of the nosecone to the specific mouse. The turquoise highlighted piece is the coil mount with different height parameters to ensure that the surface coil is as close as possible to the dorsal portion of the head. The dark purple piece is the body clamp pieces with different thicknesses to accommodate different sizes and weight classes. The dark green highlighted piece is the cover piece that holds two customizable pads that apply a slight downward pressure onto the body to reduce body movement during awake rsfMRI, there are different sizes to accommodate different locations for the pads to accommodate for different sized mice. B) A flow diagram that details the experimental paradigm from the surgical procedure to the brain dissection. C) Average framewise displacement (FD) per volume across scans that passed the data quality criteria of having at least 90% or more volumes being under the FD threshold (FD < 0.125mm). D) Average framewise displacement (FD) per volume across all scans regardless of data quality. E) Graphic that details the number of volumes that featured a FD value that exceeded the FD threshold in scans that passed the data quality criteria (number of volumes that pass FD threshold $\geq$ 90%). F) Graphic that details the number of volumes that featured a FD value that exceeded the FD threshold in all scans regardless of whether they passed or failed data quality criteria.

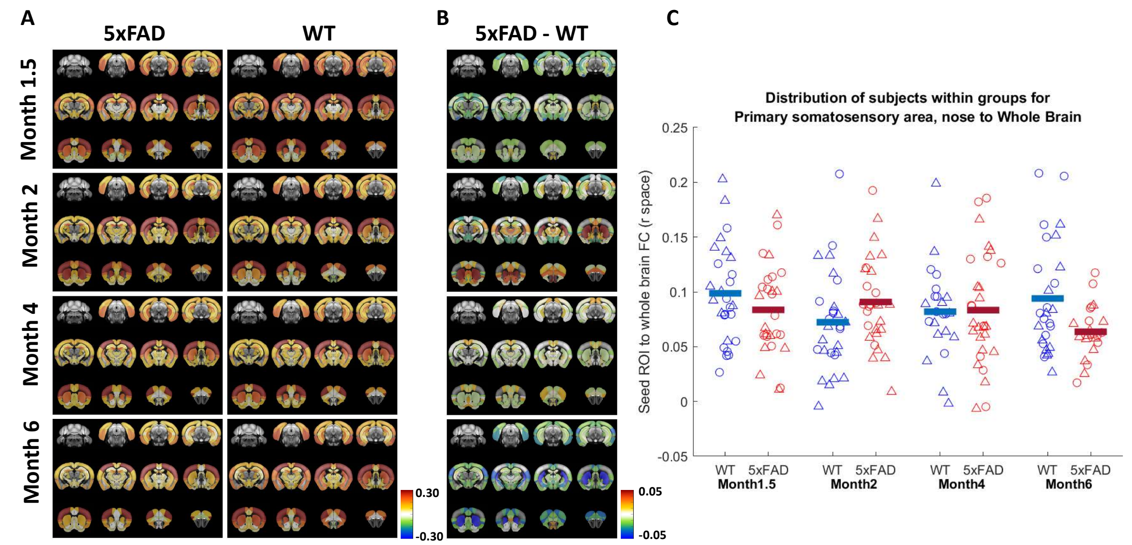

Supplementary Figure 2: Primary Somatosensory Area (S1) decouples from the whole brain over progression of Alzheimer's disease. A) Seed ROI to ROI map for both 5xFAD and WT mice for the S1 seed region for each time point. B) Difference map between 5xFAD and WT groups with the average seed ROI correlation values being used to calculate the difference values between the two genotypes controlled for timepoint. C) Distribution of subjects within each group (timepoint and genotype), with the horizontal lines denoting the group average. Sexes are coded by shape with triangles being female, and circles being male.

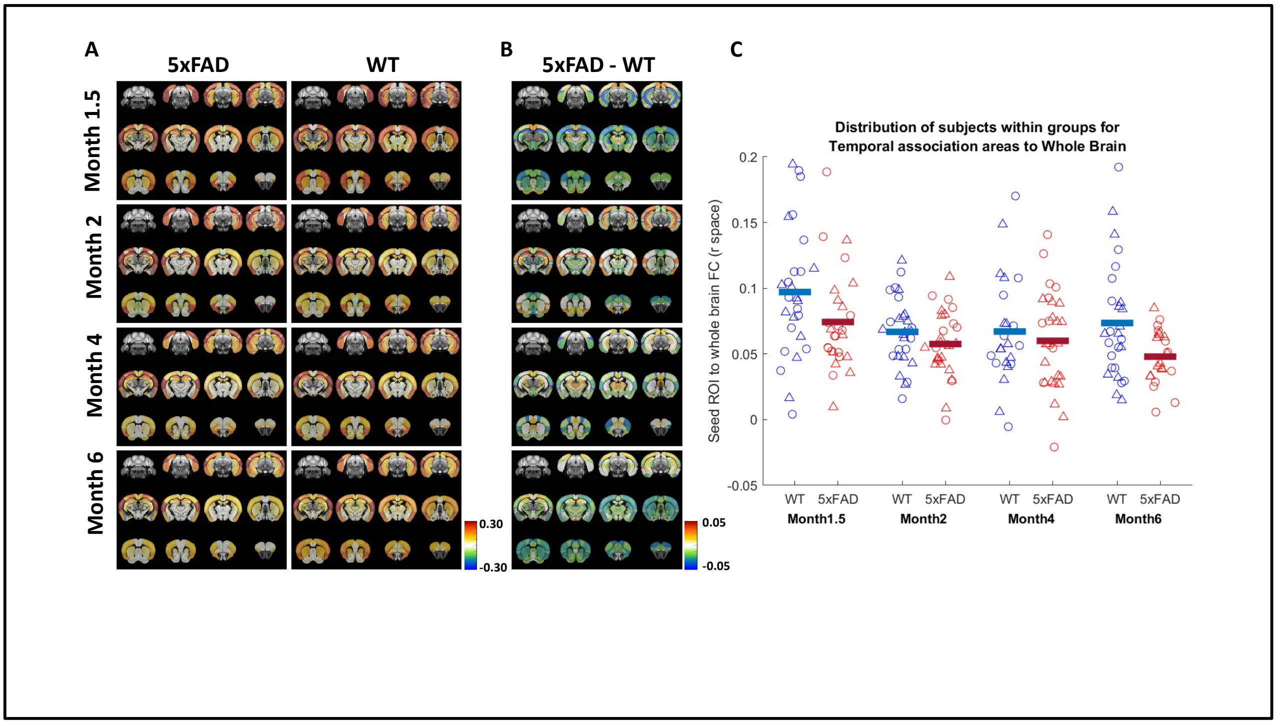

Supplementary Figure 3: Temporal Association area (TeV) decouples from the whole brain over progression of Alzheimer's disease. A) Seed ROI to ROI map for both 5xFAD and WT mice for the TeV seed region for each time point. B) Difference map between 5xFAD and WT groups with the average seed ROI correlation values being used to calculate the difference values between the two genotypes controlled for timepoint. C) Distribution of subjects within each group (timepoint and genotype), with the horizontal lines denoting the group average. Sexes are coded by shape with triangles being female, and circles being male.

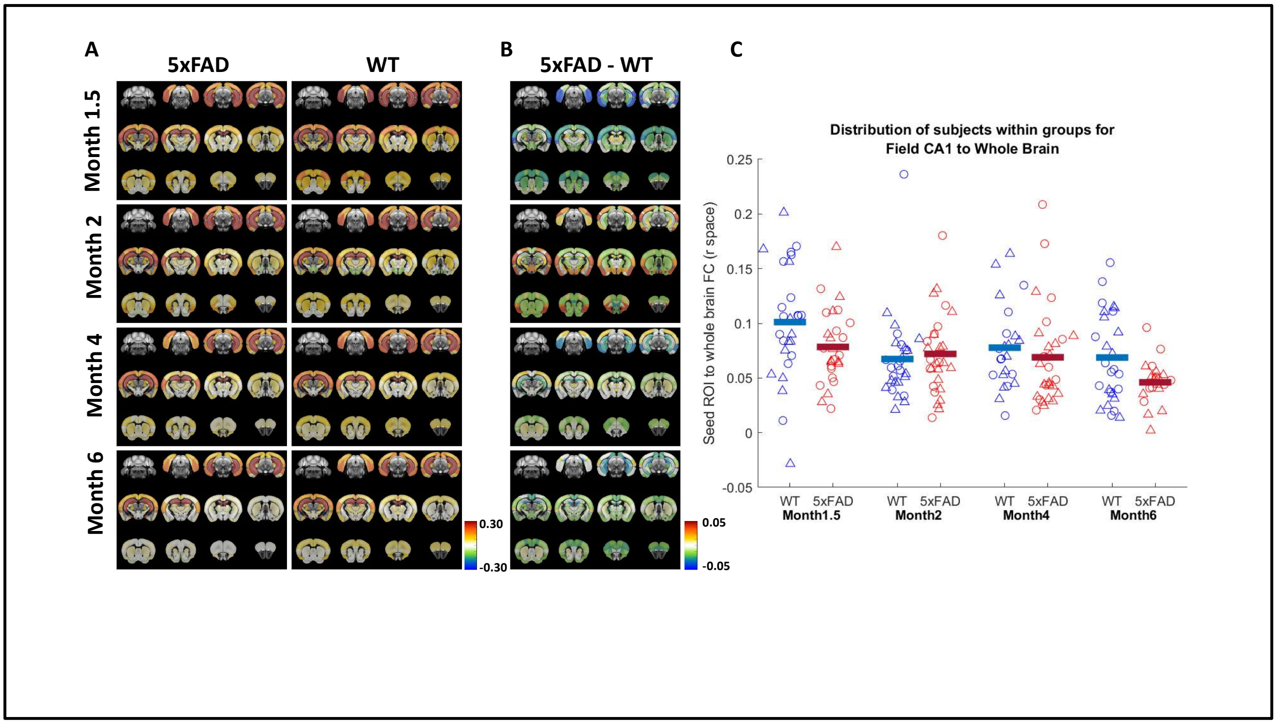

Supplementary Figure 4: Field CA1 (CA1) decouples from the whole brain over progression of Alzheimer's disease. A) Seed ROI to ROI map for both 5xFAD and WT mice for the CA1 seed region for each time point. B) Difference map between 5xFAD and WT groups with the average seed ROI correlation values being used to calculate the difference values between the two genotypes controlled for timepoint. C) Distribution of subjects within each group (timepoint and genotype), with the horizontal lines denoting the group average. Sexes are coded by shape with triangles being female, and circles being male.

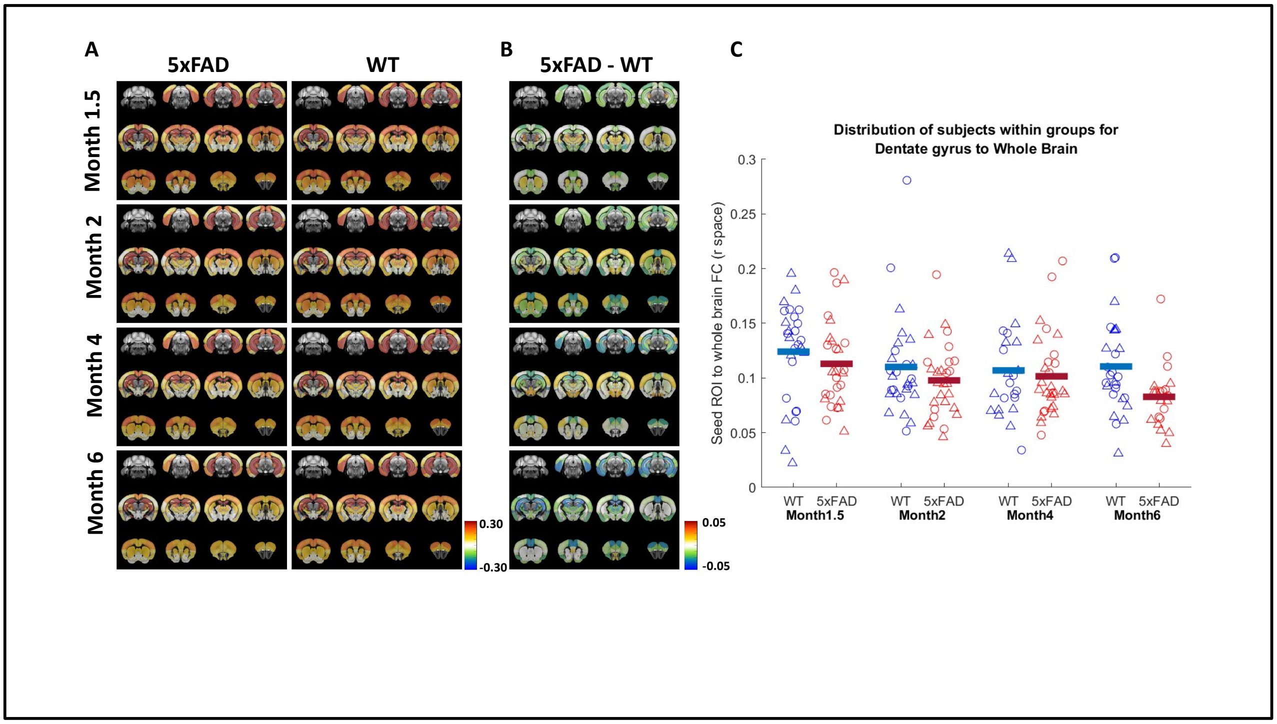

Supplementary Figure 5: Dentate Gyrus (DG) decouples from the whole brain over progression of Alzheimer's disease. A) Seed ROI to ROI map for both 5xFAD and WT mice for the DG seed region for each time point. B) Difference map between 5xFAD and WT groups with the average seed ROI correlation values being used to calculate the difference values between the two genotypes controlled for timepoint. C) Distribution of subjects within each group (timepoint and genotype), with the horizontal lines denoting the group average. Sexes are coded by shape with triangles being female, and circles being male.

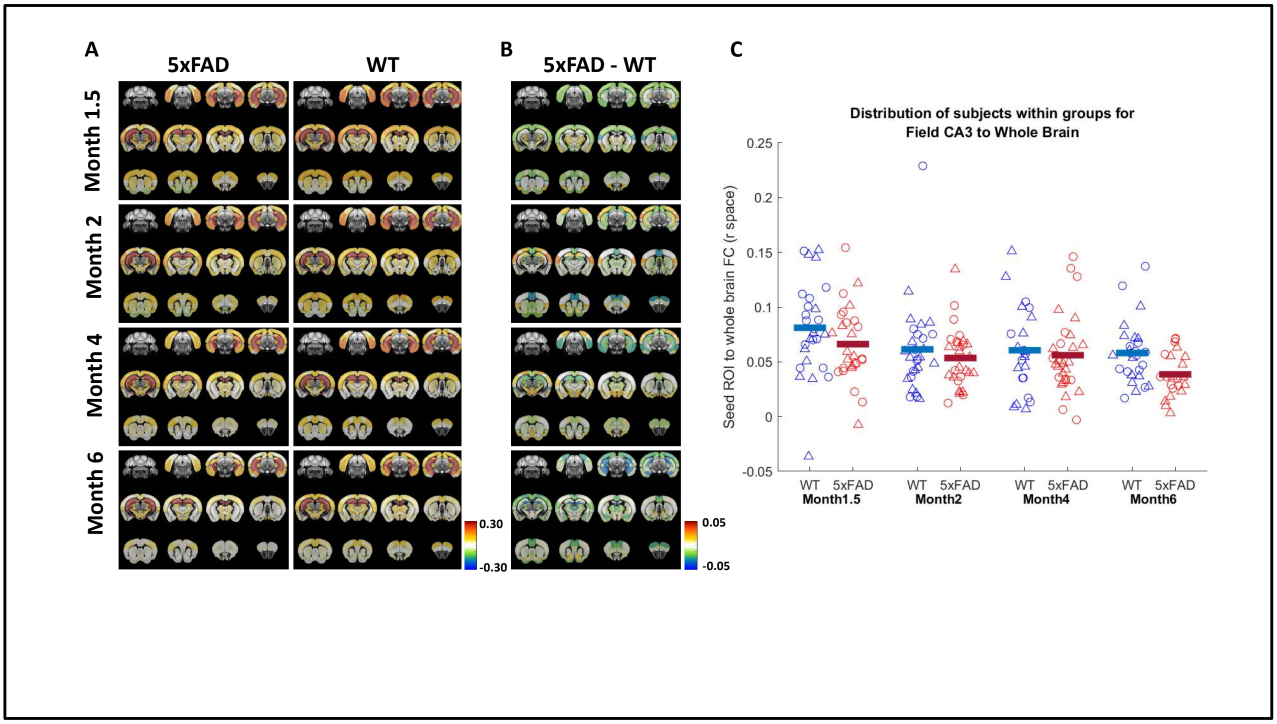

Supplementary Figure 6: Field CA3 (CA3) decouples from the whole brain over progression of Alzheimer's disease. A) Seed ROI to ROI map for both 5xFAD and WT mice for the CA3 seed region for each time point. B) Difference map between 5xFAD and WT groups with the average seed ROI correlation values being used to calculate the difference values between the two genotypes controlled for timepoint. C) Distribution of subjects within each group (timepoint and genotype), with the horizontal lines denoting the group average. Sexes are coded by shape with triangles being female, and circles being male.

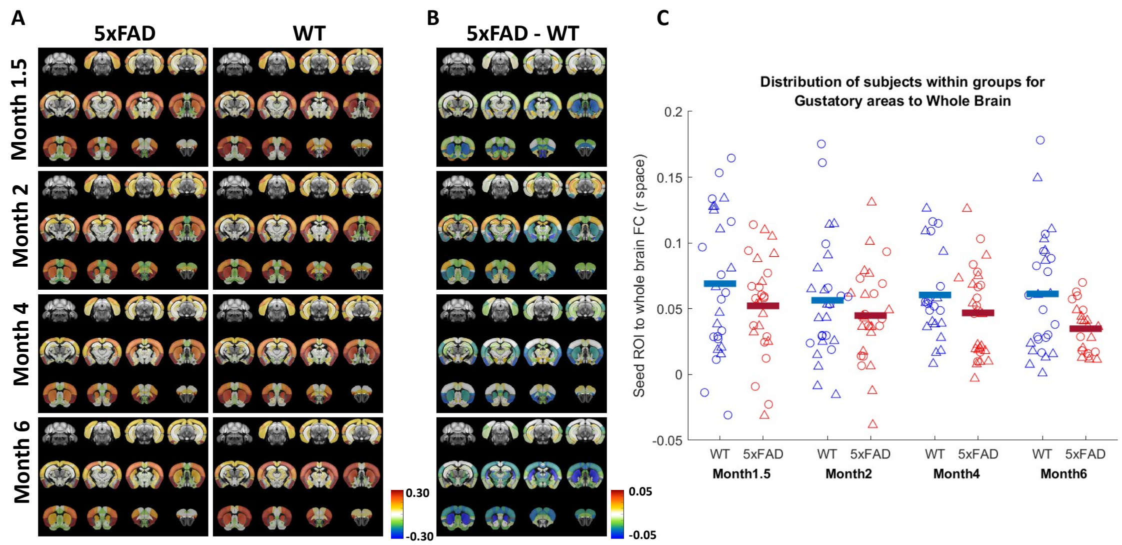

Supplementary Figure 7: Gustatory area (GC)) decouples from the whole brain over progression of Alzheimer's disease. A) Seed ROI to ROI map for both 5xFAD and WT mice for the GC seed region for each time point. B) Difference map between 5xFAD and WT groups with the average seed ROI correlation values being used to calculate the difference values between the two genotypes controlled for timepoint. C) Distribution of subjects within each group (timepoint and genotype), with the horizontal lines denoting the group average. Sexes are coded by shape with triangles being female, and circles being male.

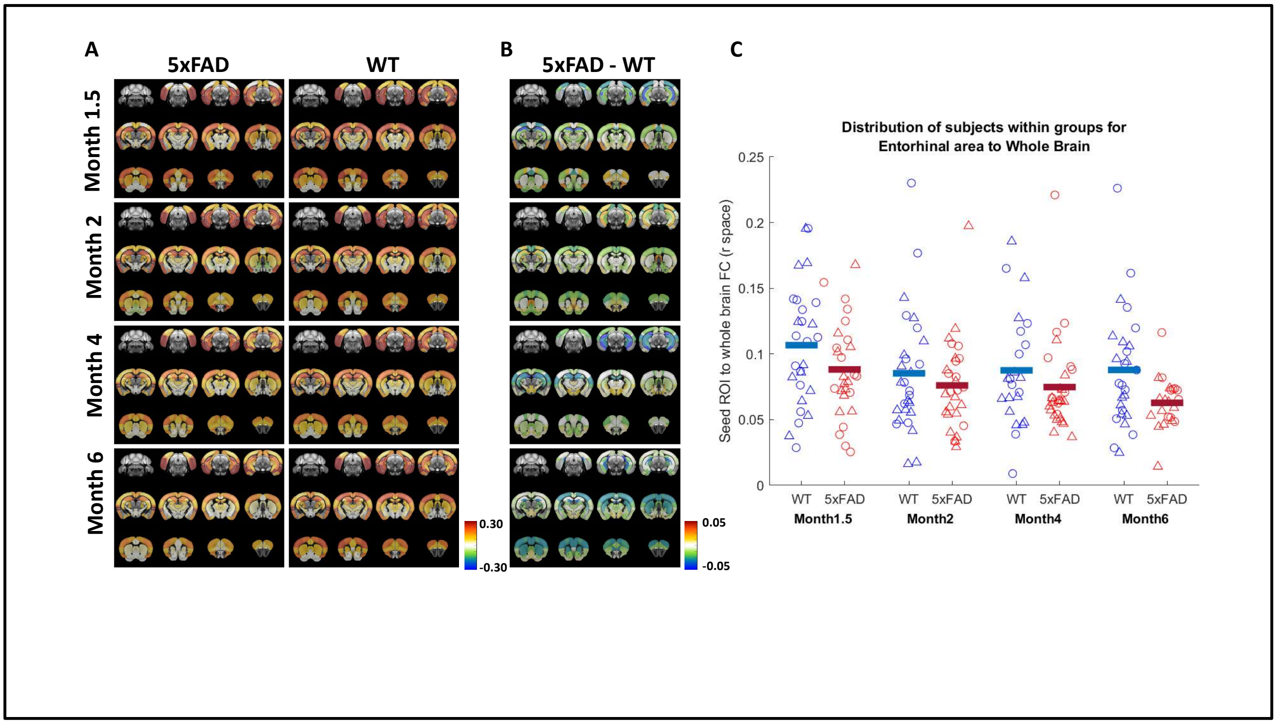

Supplementary Figure 8: Entorhinal area (EA) decouples from the whole brain over progression of Alzheimer's disease. A) Seed ROI to ROI map for both 5xFAD and WT mice for the EA seed region for each time point. B) Difference map between 5xFAD and WT groups with the average seed ROI correlation values being used to calculate the difference values between the two genotypes controlled for timepoint. C) Distribution of subjects within each group (timepoint and genotype), with the horizontal lines denoting the group average. Sexes are coded by shape with triangles being female, and circles being male.

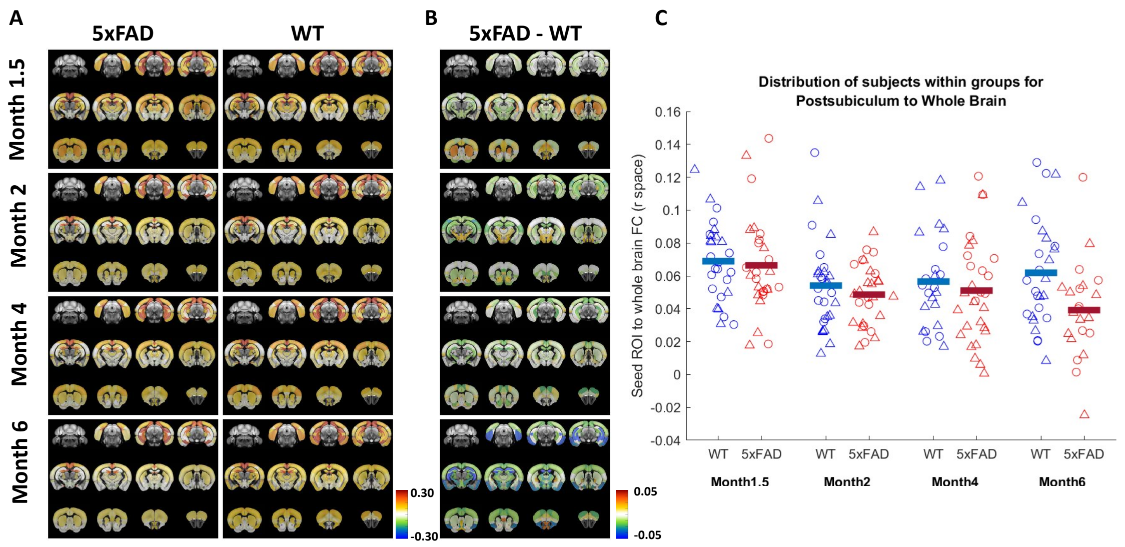

Supplementary Figure 9: Postsubiculum (PoS) decouples from the whole brain over progression of Alzheimer's disease. A) Seed ROI to ROI map for both 5xFAD and WT mice for the PoS seed region for each time point. B) Difference map between 5xFAD and WT groups with the average seed ROI correlation values being used to calculate the difference values between the two genotypes controlled for timepoint. C) Distribution of subjects within each group (timepoint and genotype), with the horizontal lines denoting the group average. Sexes are coded by shape with triangles being female, and circles being male.

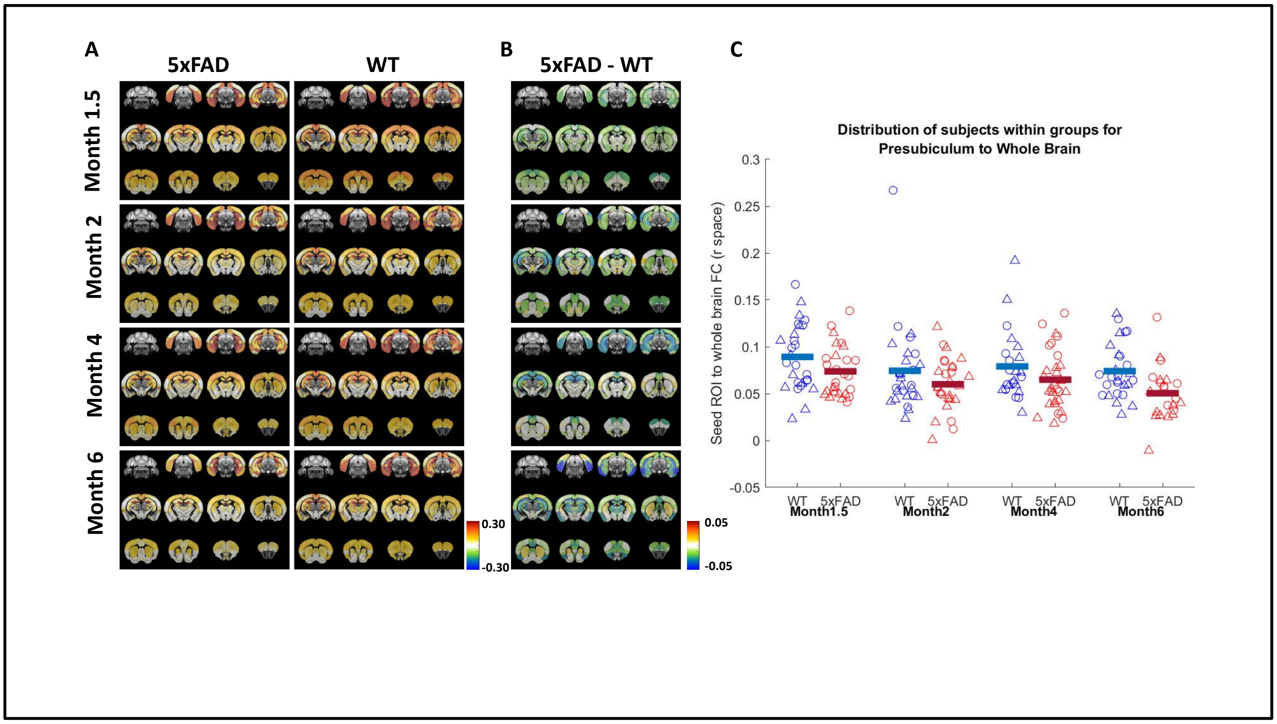

Supplementary Figure 10: Presubiculum (PrS) decouples from the whole brain over progression of Alzheimer's disease. A) Seed ROI to ROI map for both 5xFAD and WT mice for the PrS seed region for each time point. B) Difference map between 5xFAD and WT groups with the average seed ROI correlation values being used to calculate the difference values between the two genotypes controlled for timepoint. C) Distribution of subjects within each group (timepoint and genotype), with the horizontal lines denoting the group average. Sexes are coded by shape with triangles being female, and circles being male.

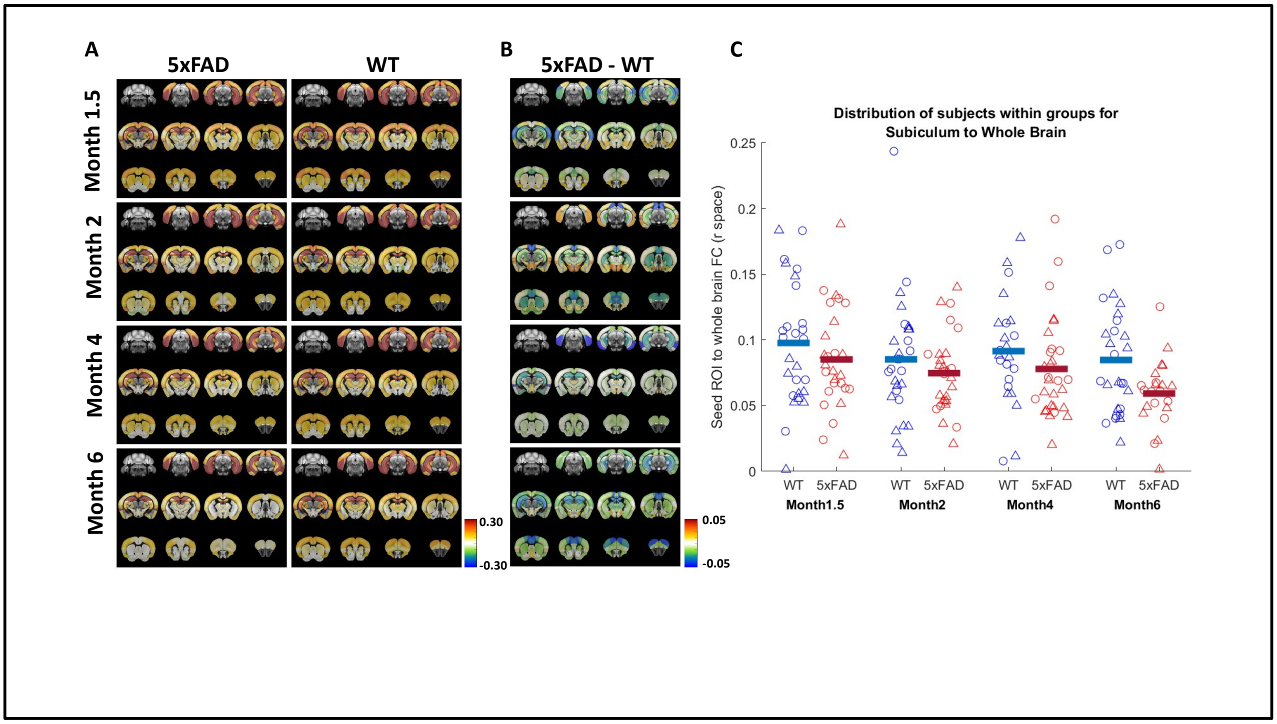

Supplementary Figure 11: Subiculum (SUB) decouples from the whole brain over progression of Alzheimer's disease. A) Seed ROI to ROI map for both 5xFAD and WT mice for the SUB seed region for each time point. B) Difference map between 5xFAD and WT groups with the average seed ROI correlation values being used to calculate the difference values between the two genotypes controlled for timepoint. C) Distribution of subjects within each group (timepoint and genotype), with the horizontal lines denoting the group average. Sexes are coded by shape with triangles being female, and circles being male.

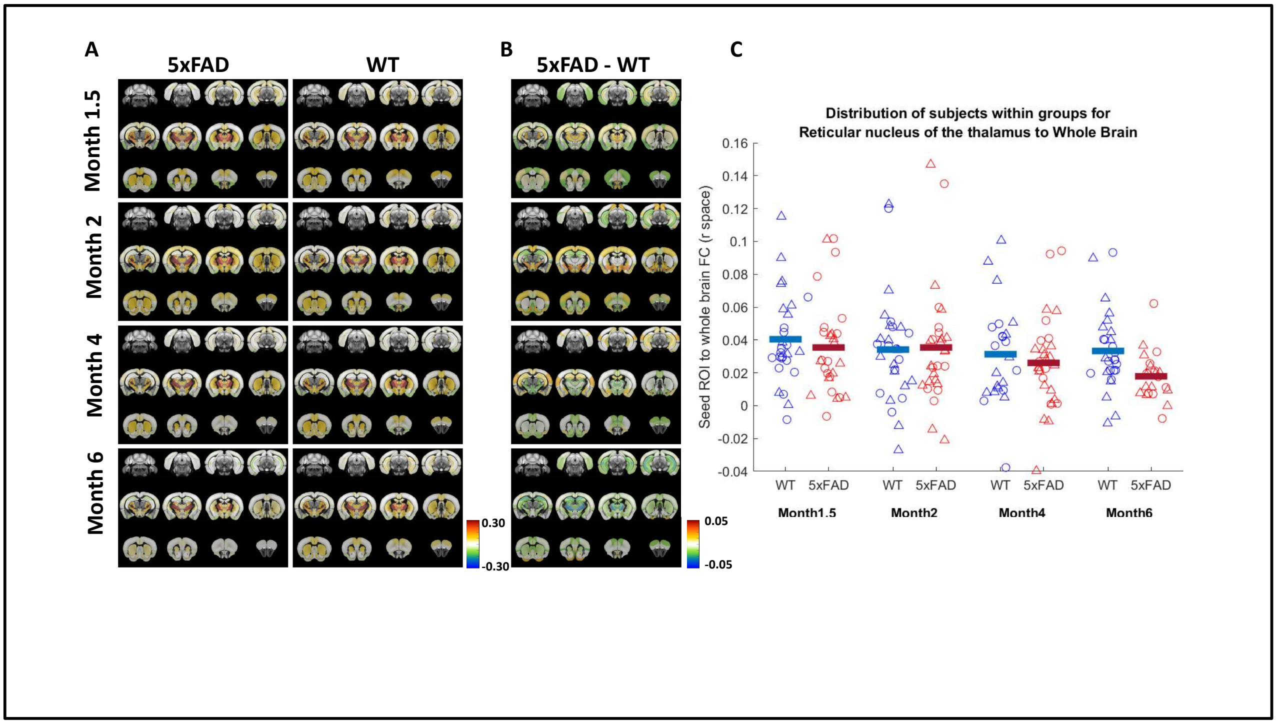

Supplementary Figure 12: Reticular nucleus of the Thalamus (TRN) decouples from the whole brain over progression of Alzheimer's disease. A) Seed ROI to ROI map for both 5xFAD and WT mice for the TRN seed region for each time point. B) Difference map between 5xFAD and WT groups with the average seed ROI correlation values being used to calculate the difference values between the two genotypes controlled for timepoint. C) Distribution of subjects within each group (timepoint and genotype), with the horizontal lines denoting the group average. Sexes are coded by shape with triangles being female, and circles being male.

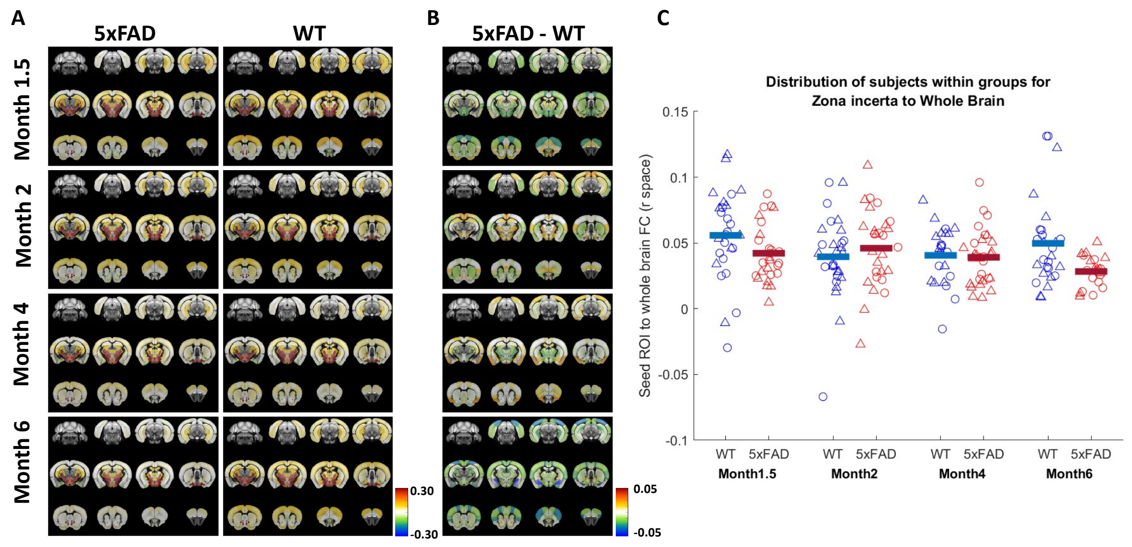

Supplementary Figure 3: Zona Incerta (ZI) decouples from the whole brain over progression of Alzheimer's disease. A) Seed ROI to ROI map for both 5xFAD and WT mice for the ZI seed region for each time point. B) Difference map between 5xFAD and WT groups with the average seed ROI correlation values being used to calculate the difference values between the two genotypes controlled for timepoint. C) Distribution of subjects within each group (timepoint and genotype), with the horizontal lines denoting the group average. Sexes are coded by shape with triangles being female, and circles being male.

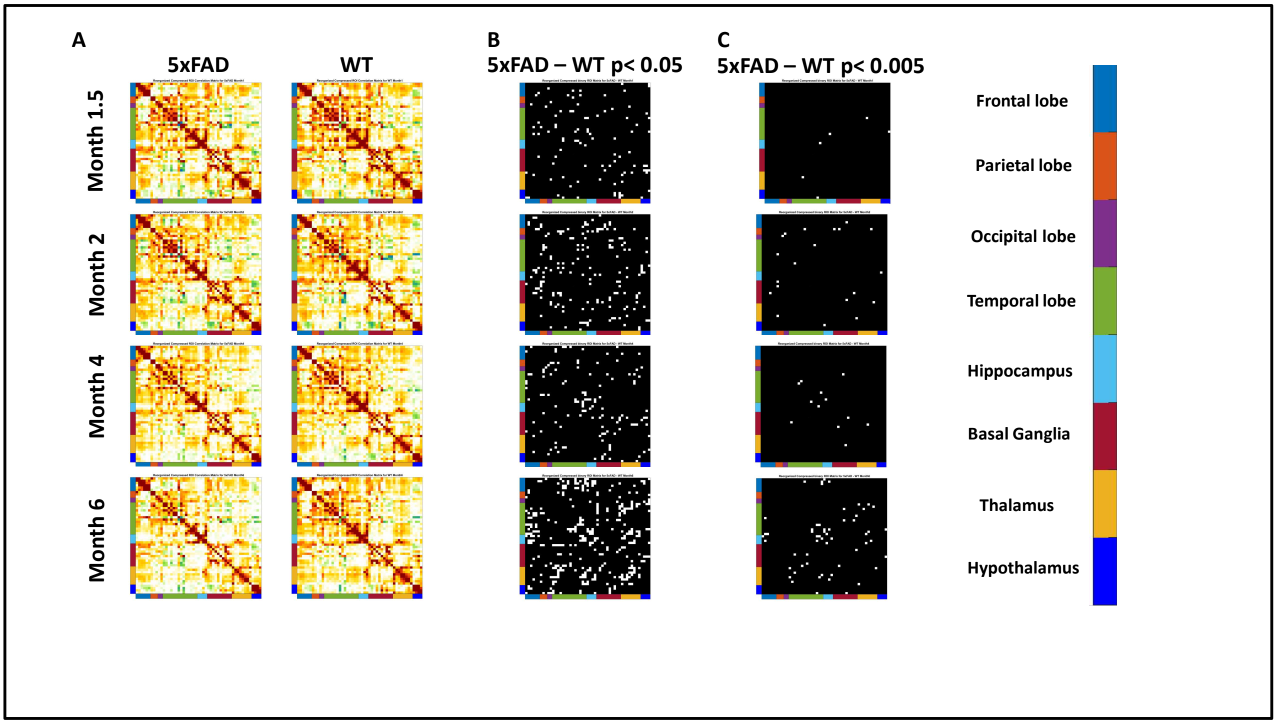

Supplementary Figure 14: Functional connectivity (FC) matrices for each group (timepoint * genotype) show gradual differences over time between genotypes. A) The group average functional connectivity matrices are displayed for each timepoint within 5xFAD, or WT genotypes. FC matrices are 51 x 51 in dimension, with 51 unique ROIs. Each ROI is grouped into anatomical regions with the legend being shown on the far right of the figure. B) Binary matrix that shows which ROI-ROI rsFC is significantly different between the 5xFAD and the WT groups at a specific time point. The statistical significance was quantified using a LME model (controlling for time) with the random effect being the scans within subject, and the fixed effects being genotype. The resulting statistical values were not corrected for multiple comparisons. C) Binary matrix that shows which ROI-ROI rsFC is significantly different between the 5xFAD and the WT groups at a specific time point. The statistical significance was quantified using a LME model (controlling for time) with the random effect being the scans within subject, and the fixed effects being genotype. The resulting statistical values were not corrected for multiple comparisons, but the threshold for statistical significance was made stricter at a threshold of 0.005, which reflects which ROI-ROI rsFC is significantly different between 5xFAD and WT groups in Figure 2B.

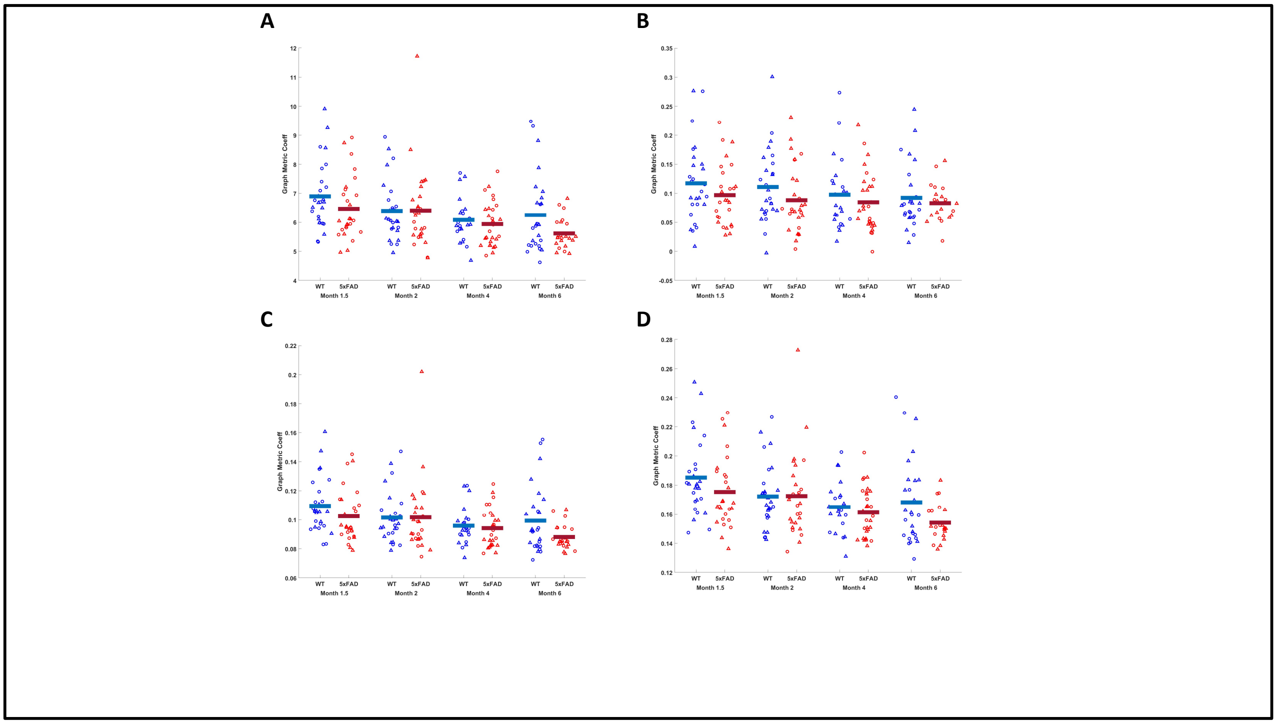

Supplementary Figure 15: Subject distribution for each group for specific network theory metrics shown in Figure 2C-F. Sexes are coded where triangles are females, and circles are males. Horizontal bars in each distribution indicates the group average. A) Subject distribution for global connectivity. B) Subject distribution for network assortativity. C) Subject distribution for global clustering D) Subject distribution for global efficiency.

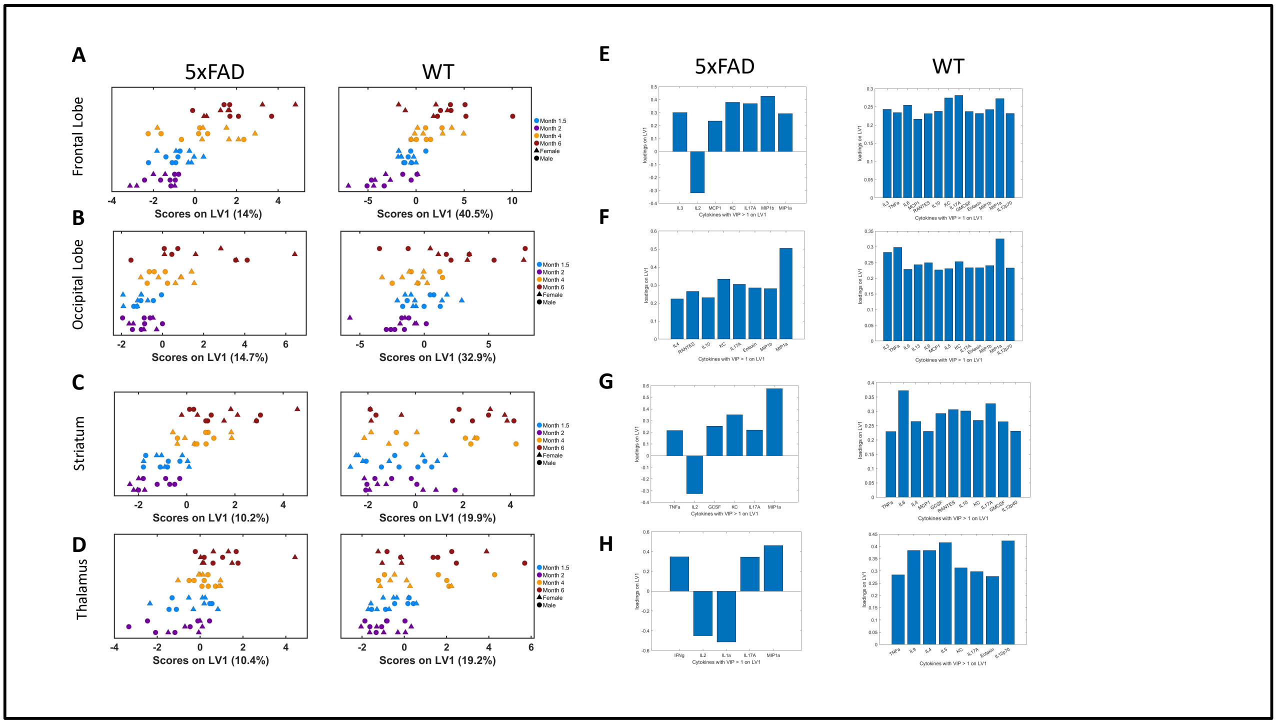

Supplementary Figure 16: Projection to latent structures regression models (PLSR) and their respective loadings for both 5xFAD and WT groups for specific brain regions. A-D: PLSR score plots for both 5xFAD and WT subjects when determining the degree of covariation between cytokine concentrations and age. A) Frontal lobe, B) Occipital lobe, C) Striatum, D) Thalamus. E-F: Loadings plots for the respective PLSR score plots. Loadings are of cytokines that had a VIP score greater than 1. The loadings corresponds to the following anatomical regions: E) Frontal lobe, F) Occipital lobe, G) Striatum, H) Thalamus.

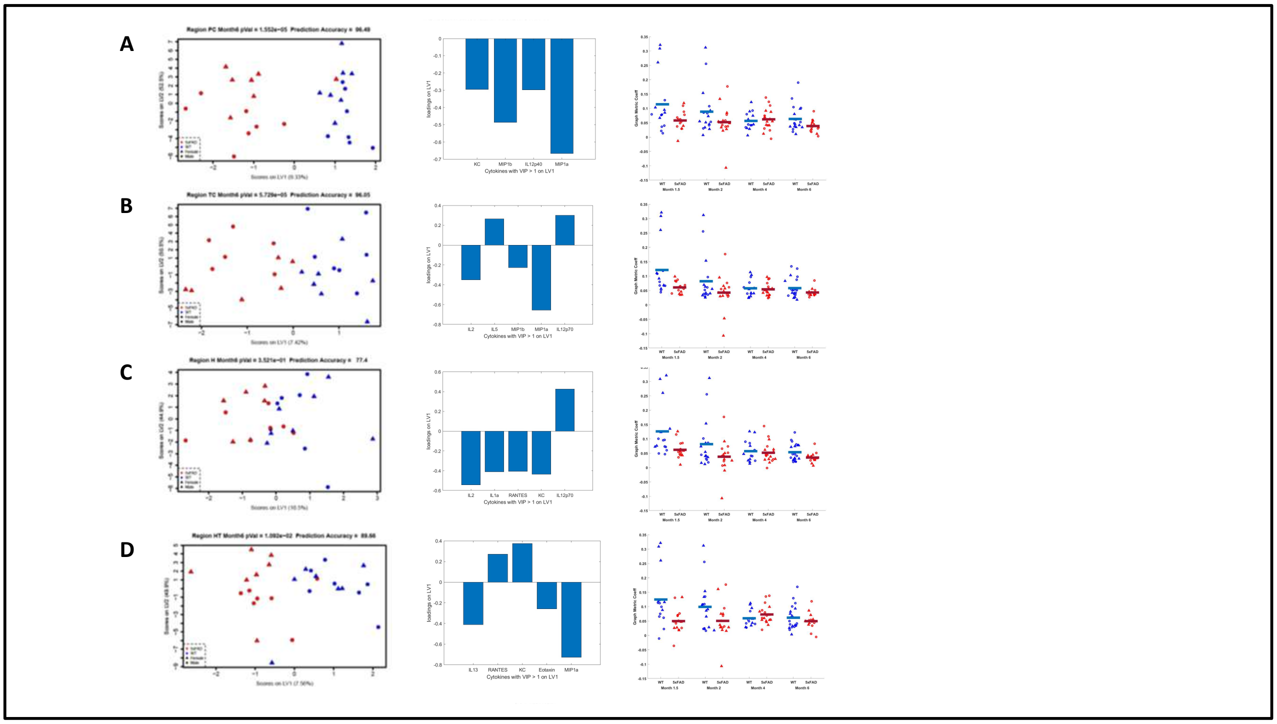

Supplementary Figure 17: Partial least squares discriminant analysis (PLSDA) score plots, loadings, and subject distribution for intersystem connectivity over disease progression. A) From left to right, the PLSDA score plots for the Parietal Cortex followed by the VIP loadings of the score plot and then the subject distribution for subject average intersystem connectivity across groups. B) From left to right, the PLSDA score plots for the Temporal Cortex followed by the VIP loadings of the score plot and then the subject distribution for subject average intersystem connectivity across groups. C) From left to right, the PLSDA score plots for the Hippocampus followed by the VIP loadings of the score plot and then the subject distribution for subject average intersystem connectivity across groups. D) From left to right, the PLSDA score plots for the Hypothalamus followed by the VIP loadings of the score plot and then the subject distribution for subject average intersystem connectivity across groups. All score plots and subject distribution scatter plots follow the same legend: red is for 5xFAD animals, blue is for WT animals. Circular points are males, and triangular points are females.

**Supplementary Table 1:** Subject meta data denoting the sex, genotype, and desired time point for animals. If the animal is designated a longitudinal animal, the time points (i.e. month 1, month 2, month 4 and month 6) indicate whether data was successfully collected with the following code: NA = not collected, collected = data was used in analysis, DISCARDED = data was removed due to motion and image quality. The inflammatory time point column indicates the time point the biological data was collected and assayed

| Sex | Genotype | Name | DOB | Time Group | Pde6B +/- | Month 1 | Month 2 | Month 4 | Month 6 | Inflammatory Time Point |
|---|---|---|---|---|---|---|---|---|---|---|
| **F** | WT | R21-5 | 4/5/2021 | Longitudinal | NA | NA | NA | Collected | Collected | NA |
| **M** | WT | R21-7 | 5/10/2021 | Month 1 | NA | Collected | NA | NA | NA | NA |
| **F** | 5xFAD | R21-8 | 5/10/2021 | Longitudinal | NA | Collected | Collected | Collected | Collected | Month6 |
| **F** | 5xFAD | R21-9 | 5/10/2021 | Longitudinal | NA | Collected | Collected | Collected | NA | NA |
| **F** | WT | R21-10 | 5/10/2021 | Longitudinal | NA | Collected | Collected | Collected | DISCARDED | DISCARDED |
| **M** | 5xFAD | R21-12 | 5/10/2021 | Longitudinal | NA | Collected | Collected | Collected | NA | Month6 |
| **M** | 5xFAD | R21-13 | 5/10/2021 | Longitudinal | NA | Collected | Collected | Collected | Collected | Month6 |
| **F** | 5xFAD | R21-14 | 5/10/2021 | Longitudinal | NA | Collected | Collected | Collected | Collected | Month6 |
| **F** | WT | R21-15 | 7/6/2021 | Longitudinal | NA | Collected | Collected | DISCARDED | DISCARDED | DISCARDED |
| **F** | WT | R21-16 | 7/6/2021 | Longitudinal | NA | Motion Issues | Motion Issues | Collected | Collected | Month6 |
| **F** | 5xFAD | R21-17 | 7/6/2021 | Longitudinal | NA | Collected | Collected | Collected | Collected | Month6 |
| **F** | WT | R21-18 | 7/6/2021 | Longitudinal | NA | NA | Collected | Collected | Collected | Month6 |
| **M** | WT | R21-19 | 7/6/2021 | Longitudinal | NA | Collected | Collected | NA | NA | NA |

| | | | | | | | | | | |
|---|---|---|---|---|---|---|---|---|---|---|
| **M** | WT | R21-22 | 7/6/2021 | Longitudinal | NA | Collected | Collected | DISCARDED | DISCARDED | DISCARDED |
| **F** | 5xFAD | R21-23 | 12/1/2021 | Longitudinal | NA | Collected | Collected | Collected | DISCARDED | DISCARDED |
| **F** | 5xFAD | R21-26 | 12/1/2021 | Longitudinal | NA | Collected | Collected | Collected | Collected | DISCARDED |
| **M** | WT | R21-27 | 1/4/2022 | Longitudinal | -/? | Collected | Collected | Collected | Collected | NA |
| **M** | 5xFAD | R21-28 | 1/4/2022 | Longitudinal | -/? | Collected | Collected | DISCARDED | DISCARDED | DISCARDED |
| **M** | WT | R21-29 | 1/4/2022 | Longitudinal | -/? | Collected | Motion Issues | Collected | Collected | Month6 |
| **M** | 5xFAD | R21-31 | 1/4/2022 | Longitudinal | -/? | Collected | Collected | DISCARDED | DISCARDED | DISCARDED |
| **F** | WT | R21-32 | 1/4/2022 | Longitudinal | -/? | Collected | Collected | Collected | Collected | Month6 |
| **M** | 5xFAD | R21-33 | 1/4/2022 | Longitudinal | -/? | Collected | Collected | Collected | Collected | Month6 |
| **M** | WT | R21-35 | 3/21/2022 | Month 6 | -/? | NA | NA | NA | Collected | Month6 |
| **F** | WT | R21-36 | 3/21/2022 | Longitudinal | -/? | Collected | Collected | DISCARDED | DISCARDED | DISCARDED |
| **M** | WT | R21-38 | 3/21/2022 | Longitudinal | -/? | Collected | Collected | Collected | Collected | Month6 |
| **F** | 5xFAD | R21-39 | 3/21/2022 | Longitudinal | -/? | Collected | Motion Issues | Collected | DISCARDED | DISCARDED |
| **M** | WT | R21-40 | 3/21/2022 | Longitudinal | -/? | Collected | Collected | Collected | NA | NA |
| **F** | WT | R21-41 | 3/21/2022 | Longitudinal | -/? | Collected | Collected | DISCARDED | DISCARDED | DISCARDED |

| | | | | | | | | | | |
|---|---|---|---|---|---|---|---|---|---|---|
| **M** | WT | R21-42 | 3/21/2022 | Month 6 | -/? | NA | NA | NA | Collected | Month6 |
| **M** | WT | R21-44 | 3/21/2022 | Month 6 | -/? | NA | NA | NA | Collected | Month6 |
| **M** | WT | R21-45 | 5/11/2022 | Month 4 | -/? | NA | NA | Collected | NA | Month4 |
| **F** | 5xFAD | R21-46 | 5/11/2022 | Month 4 | -/? | NA | NA | Collected | NA | Month4 |
| **F** | WT | R21-47 | 5/11/2022 | Month 4 | -/? | NA | NA | Collected | NA | Month4 |
| **F** | 5xFAD | R21-48 | 5/11/2022 | Month 4 | -/? | NA | NA | Collected | NA | Month4 |
| **F** | 5xFAD | R21-49 | 5/11/2022 | Month 4 | -/? | NA | NA | Collected | NA | Month4 |
| **M** | WT | R21-51 | 5/11/2022 | Month 4 | -/? | NA | NA | Collected | NA | Month4 |
| **M** | 5xFAD | R21-52 | 5/11/2022 | Month 4 | -/? | NA | NA | Collected | NA | Month4 |
| **M** | WT | R21-53 | 5/11/2022 | Month 4 | -/? | NA | NA | Collected | NA | Month4 |
| **F** | 5xFAD | R21-54 | 5/11/2022 | Month 4 | -/? | NA | NA | Collected | NA | Month4 |
| **F** | 5xFAD | R21-55 | 5/11/2022 | Month 4 | -/? | NA | NA | Collected | NA | Month4 |
| **F** | WT | R21-56 | 5/11/2022 | Month 4 | -/? | NA | NA | Collected | NA | Month4 |
| **F** | WT | R21-57 | 5/11/2022 | Month 4 | -/? | NA | NA | Collected | NA | Month4 |
| **F** | 5xFAD | R21-58 | 5/13/2022 | Longitudinal | -/? | NA | Collected | Collected | Collected | Month6 |
| **F** | WT | R21-59 | 5/13/2022 | Month 6 | -/? | NA | NA | NA | Collected | Month6 |
| **M** | WT | R21-61 | 5/13/2022 | Month 6 | -/? | NA | NA | NA | Collected | Month6 |
| **M** | 5xFAD | R21-62 | 5/13/2022 | Month 6 | -/? | NA | NA | NA | Collected | Month6 |

| | | | | | | | | | | |
|---|---|---|---|---|---|---|---|---|---|---|
| **F** | 5xFAD | R21-72 | 6/3/2022 | Month 4 | -/? | NA | NA | Collected | NA | Month4 |
| **F** | WT | R21-73 | 6/3/2022 | Month 4 | -/? | NA | NA | Collected | NA | Month4 |
| **M** | 5xFAD | R21-74 | 6/3/2022 | Month 4 | -/? | NA | NA | Collected | NA | Month4 |
| **M** | WT | R21-75 | 6/3/2022 | Month 4 | -/? | NA | NA | Collected | NA | Month4 |
| **F** | 5xFAD | R21-76 | 7/19/2022 | Month 1 | -/? | Collected | NA | NA | NA | Month1 |
| **M** | WT | R21-77 | 7/19/2022 | Month 1 | -/? | Collected | NA | NA | NA | Month1 |
| **M** | 5xFAD | R21-78 | 7/19/2022 | Month 1 | -/? | Collected | NA | NA | NA | Month1 |
| **M** | 5xFAD | R21-80 | 7/19/2022 | Month 1 | -/? | Collected | NA | NA | NA | Month1 |
| **F** | 5xFAD | R21-81 | 7/19/2022 | Month 1 | -/? | Collected | NA | NA | NA | Month1 |
| **M** | 5xFAD | R21-82 | 7/19/2022 | Month 1 | -/? | Collected | NA | NA | NA | Month1 |
| **F** | 5xFAD | R21-83 | 7/19/2022 | Month 1 | -/? | Collected | NA | NA | NA | Month1 |
| **M** | 5xFAD | R21-84 | 7/19/2022 | Month 2 | -/? | NA | Collected | NA | NA | Month2 |
| **F** | 5xFAD | R21-85 | 7/19/2022 | Month 2 | -/? | NA | Collected | NA | NA | Month2 |
| **M** | 5xFAD | R21-86 | 7/19/2022 | Month 2 | -/? | NA | Collected | NA | NA | Month2 |
| **F** | 5xFAD | R21-87 | 7/19/2022 | Month 2 | -/? | NA | Collected | NA | NA | Month2 |
| **M** | WT | R21-88 | 7/19/2022 | Month 2 | -/? | NA | Collected | NA | NA | Month2 |
| **M** | WT | R21-89 | 7/19/2022 | Month 2 | -/? | NA | Collected | NA | NA | Month2 |
| **M** | 5xFAD | R21-90 | 7/19/2022 | Month 1 | -/? | Collected | NA | NA | NA | Month1 |

| | | | | | | | | | | |
|---|---|---|---|---|---|---|---|---|---|---|
| **M** | 5xFAD | R21-91 | 7/19/2022 | Month 2 | -/? | NA | Collected | NA | NA | Month2 |
| **F** | WT | R21-92 | 7/19/2022 | Month 2 | -/? | NA | Collected | NA | NA | Month2 |
| **F** | 5xFAD | R21-93 | 7/19/2022 | Month 2 | -/? | NA | Collected | NA | NA | Month2 |
| **M** | WT | R21-94 | 9/3/2022 | Month 6 | -/? | NA | NA | NA | Collected | Month6 |
| **F** | WT | R21-96 | 9/3/2022 | Month 6 | -/? | NA | NA | NA | Collected | Month6 |
| **M** | WT | R21-97 | 9/3/2022 | Month 6 | -/? | NA | NA | NA | Collected | Month6 |
| **M** | 5xFAD | R21-100 | 9/3/2022 | Month 6 | -/? | NA | NA | NA | Collected | Month6 |
| **M** | 5xFAD | R21-101 | 9/3/2022 | Month 6 | -/? | NA | NA | NA | Collected | Month6 |
| **M** | 5xFAD | R21-103 | 9/4/2022 | Month 6 | -/? | NA | NA | NA | Collected | Month6 |
| **M** | 5xFAD | R21-105 | 10/23/2022 | Month 4 | -/? | NA | NA | Collected | NA | Month4 |
| **M** | 5xFAD | R21-106 | 10/23/2022 | Month 4 | -/? | NA | NA | Collected | NA | Month4 |
| **M** | 5xFAD | R21-107 | 10/23/2022 | Month 4 | -/? | NA | NA | Collected | NA | Month4 |
| **M** | WT | R21-108 | 10/23/2022 | Month 6 | -/? | NA | NA | NA | Collected | Month6 |
| **M** | 5xFAD | R21-110 | 10/23/2022 | Month 6 | -/? | NA | NA | NA | Collected | Month6 |

| | | | | | | | | | | |
|---|---|---|---|---|---|---|---|---|---|---|
| **F** | 5xFAD | R21-112 | 10/23/2022 | Month 6 | -/? | NA | NA | NA | Collected | Month6 |
| **M** | WT | R21-113 | 10/23/2022 | Month 6 | -/? | NA | NA | NA | Collected | Month6 |
| **F** | WT | R21-114 | 10/23/2022 | Month 6 | -/? | NA | NA | NA | Collected | Month6 |
| **F** | 5xFAD | R21-116 | 11/11/2022 | Month 2 | -/? | NA | Collected | NA | NA | Month2 |
| **F** | 5xFAD | R21-117 | 11/11/2022 | Month 2 | -/? | NA | Collected | NA | NA | Month2 |
| **M** | WT | R21-119 | 11/11/2022 | Month 2 | -/? | NA | Collected | NA | NA | Month2 |
| **M** | WT | R21-120 | 11/11/2022 | Month 2 | -/? | NA | Collected | NA | NA | Month2 |
| **F** | 5xFAD | R21-121 | 11/16/2022 | Month 1 | -/? | Collected | NA | NA | NA | Month1 |
| **F** | WT | R21-122 | 11/16/2022 | Month 1 | -/? | Collected | NA | NA | NA | Month1 |
| **F** | WT | R21-124 | 11/16/2022 | Month 1 | -/? | Collected | NA | NA | NA | Month1 |
| **M** | WT | R21-125 | 11/16/2022 | Month 1 | -/? | Collected | NA | NA | NA | Month1 |
| **F** | 5xFAD | R21-126 | 11/16/2022 | Month 1 | -/? | Collected | NA | NA | NA | Month1 |
| **M** | 5xFAD | R21-127 | 11/16/2022 | Month 1 | -/? | Collected | NA | NA | NA | Month1 |

| | | | | | | | | | | |
|---|---|---|---|---|---|---|---|---|---|---|
| **M** | 5xFAD | R21-129 | 11/16/2022 | Month 1 | -/? | Collected | NA | NA | NA | Month1 |
| **M** | 5xFAD | R21-130 | 9/25/2022 | Month 6 | -/? | NA | NA | NA | Collected | Month6 |
| **M** | 5xFAD | R21-131 | 9/25/2022 | Month 6 | -/? | NA | NA | NA | Collected | Month6 |
| **M** | 5xFAD | R21-132 | 9/25/2022 | Month 6 | -/? | NA | NA | NA | Collected | Month6 |
| **F** | 5xFAD | R21-133 | 9/25/2022 | Month 6 | -/? | NA | NA | NA | Collected | Month6 |
| **F** | WT | R21-134 | 9/25/2022 | Month 6 | -/? | NA | NA | NA | Collected | Month6 |
| **M** | WT | R21-135 | 9/25/2022 | Month 6 | -/? | NA | NA | NA | Collected | Month6 |
| **F** | 5xFAD | R21-136 | 9/25/2022 | Month 6 | -/? | NA | NA | NA | Collected | Month6 |
| **M** | WT | R21-137 | 9/25/2022 | Month 6 | -/? | NA | NA | NA | Collected | Month6 |
| **M** | WT | R21-138 | 9/25/2022 | Month 6 | -/? | NA | NA | NA | Collected | Month6 |
| **F** | WT | R21-139 | 9/25/2022 | Month 6 | -/? | NA | NA | NA | Collected | Month6 |
| **F** | WT | R21-140 | 12/6/2022 | Month 2 | -/- | NA | Collected | NA | NA | Month2 |
| **F** | WT | R21-141 | 12/6/2022 | Month 2 | -/- | NA | Collected | NA | NA | Month2 |

| | | | | | | | | | | |
|---|---|---|---|---|---|---|---|---|---|---|
| **F** | 5xFAD | R21-142 | 12/6/2022 | Month 2 | -/- | NA | Collected | NA | NA | Month2 |
| **F** | WT | R21-143 | 12/6/2022 | Month 2 | -/- | NA | Collected | NA | NA | Month2 |
| **M** | WT | R21-144 | 12/6/2022 | Month 2 | -/- | NA | Collected | NA | NA | Month2 |
| **M** | WT | R21-145 | 12/6/2022 | Month 2 | -/- | NA | Collected | NA | NA | Month2 |
| **F** | WT | R21-146 | 12/6/2022 | Month 2 | -/- | NA | Collected | NA | NA | Month2 |
| **M** | 5xFAD | R21-148 | 12/6/2022 | Month 2 | -/- | NA | Collected | NA | NA | Month2 |
| **F** | WT | R21-149 | 12/14/2022 | Month 1 | -/? | Collected | NA | NA | NA | Month1 |
| **F** | WT | R21-150 | 12/14/2022 | Month 1 | -/? | Collected | NA | NA | NA | Month1 |
| **M** | WT | R21-151 | 12/14/2022 | Month 1 | -/? | Collected | NA | NA | NA | Month1 |
| **F** | WT | R21-152 | 12/14/2022 | Month 1 | -/? | Collected | NA | NA | NA | Month1 |
| **M** | WT | R21-153 | 12/14/2022 | Month 1 | -/? | Collected | NA | NA | NA | Month1 |
| **M** | WT | R21-154 | 12/14/2022 | Month 1 | -/? | Collected | NA | NA | NA | Month1 |
| **M** | 5xFAD | R21-155 | 12/14/2022 | Month 1 | -/? | Collected | NA | NA | NA | Month1 |

| | | | | | | | | | | |
|---|---|---|---|---|---|---|---|---|---|---|
| **M** | WT | R21-156 | 12/14/2022 | Month 1 | -/? | Collected | NA | NA | NA | Month1 |
| **F** | WT | R21-157 | 2/2/2023 | Month 2 | -/? | NA | Collected | NA | NA | Month2 |
| **M** | 5xFAD | R21-158 | 2/2/2023 | Month 2 | -/? | NA | Collected | NA | NA | Month2 |
| **M** | 5xFAD | R21-159 | 2/2/2023 | Month 2 | -/? | NA | Collected | NA | NA | Month2 |
| **M** | WT | R21-160 | 2/2/2023 | Month 2 | -/? | NA | Collected | NA | NA | Month2 |
| **F** | 5xFAD | R21-161 | 2/2/2023 | Month 2 | -/? | NA | Collected | NA | NA | Month2 |
| **F** | 5xFAD | R21-162 | 2/2/2023 | Month 2 | -/? | NA | Collected | NA | NA | Month2 |
| **F** | 5xFAD | R21-163 | 2/2/2023 | Month 2 | -/? | NA | Collected | NA | NA | Month2 |
| **F** | 5xFAD | R21-164 | 2/2/2023 | Month 2 | -/? | NA | Collected | NA | NA | Month2 |
| **F** | 5xFAD | R21-165 | 1/20/2023 | Month 4 | -/? | NA | NA | Collected | NA | Month4 |
| **F** | WT | R21-166 | 1/20/2023 | Month 4 | -/? | NA | NA | Collected | NA | Month4 |
| **F** | 5xFAD | R21-167 | 1/20/2023 | Month 4 | -/? | NA | NA | Collected | NA | Month4 |
| **M** | 5xFAD | R21-168 | 1/20/2023 | Month 4 | -/? | NA | NA | Collected | NA | Month4 |

| | | | | | | | | | | |
|---|---|---|---|---|---|---|---|---|---|---|
| **M** | WT | R21-169 | 1/20/2023 | Month 4 | -/? | NA | NA | Collected | NA | Month4 |
| **F** | 5xFAD | R21-170 | 1/6/2023 | Month 6 | -/? | NA | NA | NA | Collected | Month6 |
| **F** | WT | R21-171 | 1/6/2023 | Month 6 | -/? | NA | NA | NA | Collected | Month6 |
| **F** | 5xFAD | R21-172 | 1/6/2023 | Month 6 | -/? | NA | NA | NA | Collected | Month6 |
| **F** | 5xFAD | R21-173 | 12/30/2022 | Month 6 | -/? | NA | NA | NA | Collected | Month6 |
| **F** | 5xFAD | R21-174 | 12/30/2022 | Month 6 | -/? | NA | NA | NA | Collected | Month6 |
| **M** | 5xFAD | R21-175 | 2/10/2023 | Month 4 | -/? | NA | NA | Collected | NA | Month4 |
| **F** | WT | R21-176 | 2/10/2023 | Month 4 | -/? | NA | NA | Collected | NA | Month4 |
| **F** | WT | R21-177 | 2/10/2023 | Month 4 | -/? | NA | NA | Collected | NA | Month4 |
| **F** | WT | R21-178 | 2/10/2023 | Month 4 | -/? | NA | NA | Collected | NA | Month4 |
| **F** | 5xFAD | R21-179 | 2/10/2023 | Month 4 | -/? | NA | NA | Collected | NA | Month4 |
| **F** | 5xFAD | R21-180 | 2/10/2023 | Month 4 | -/? | NA | NA | Collected | NA | Month4 |
| **M** | 5xFAD | R21-181 | 3/6/2023 | Month 4 | -/? | NA | NA | Collected | NA | Month4 |

| | | | | | | | | | | |
|---|---|---|---|---|---|---|---|---|---|---|
| **M** | WT | R21-182 | 3/6/2023 | Month 4 | -/? | NA | NA | Collected | NA | Month4 |
| **F** | WT | R21-183 | 1/3/2023 | Month 6 | -/? | NA | NA | NA | Collected | Month6 |
| **M** | 5xFAD | R21-185 | 5/15/2023 | Month 1 | -/? | Collected | NA | NA | NA | Month1 |
| **M** | 5xFAD | R21-186 | 5/15/2023 | Month 1 | -/? | Collected | NA | NA | NA | Month1 |
| **M** | WT | R21-187 | 5/15/2023 | Month 1 | -/? | Collected | NA | NA | NA | Month1 |
| **M** | WT | R21-188 | 5/15/2023 | Month 1 | -/? | Collected | NA | NA | NA | Month1 |
| **M** | 5xFAD | R21-189 | 5/15/2023 | Month 1 | -/? | Collected | NA | NA | NA | Month1 |
| **F** | WT | R21-191 | 5/15/2023 | Month 1 | -/? | Collected | NA | NA | NA | Month1 |
| **F** | WT | R21-192 | 5/15/2023 | Month 1 | -/? | Collected | NA | NA | NA | Month1 |
| **F** | WT | R21-193 | 1/3/2023 | Month 2 | -/? | NA | Collected | NA | NA | Month2 |
| **F** | WT | R21-194 | 1/3/2023 | Month 2 | -/? | NA | Collected | NA | NA | Month2 |
| **F** | WT | R21-195 | 5/10/2023 | Month 2 | -/? | NA | Collected | NA | NA | Month2 |
| **F** | WT | R21-196 | 5/10/2023 | Month 2 | -/? | NA | Collected | NA | NA | Month2 |

| | | | | | | | | | | |
|---|---|---|---|---|---|---|---|---|---|---|
| **F** | WT | R21-197 | 5/10/2023 | Month 2 | -/? | NA | Collected | NA | NA | Month2 |
| **F** | WT | R21-198 | 5/10/2023 | Month 2 | -/? | NA | Collected | NA | NA | Month2 |

**Supplementary Table 2:** Table listing the designations of the different ROIs and the system they are assigned.

| Regions of Interest | Assigned System |
|---|---|
| **Primary motor area** | Frontal Lobe |
| **Secondary motor area** | Frontal Lobe |
| **Anterior cingulate area** | Frontal Lobe |
| **Prelimbic area** | Frontal Lobe |
| **Infralimbic area** | Frontal Lobe |
| **Orbital area** | Frontal Lobe |
| **Primary somatosensory area, nose** | Parietal Lobe |
| **Supplemental somatosensory area** | Parietal Lobe |
| **Posterior parietal association areas** | Parietal Lobe |
| **Visual areas** | Occipital Lobe |
| **Retrosplenial area** | Occipital Lobe |
| **Gustatory areas** | Temporal Lobe |
| **Visceral area** | Temporal Lobe |
| **Auditory areas** | Temporal Lobe |
| **Agranular insular area** | Temporal Lobe |
| **Temporal association areas** | Temporal Lobe |
| **Ectorhinal area** | Temporal Lobe |
| **Taenia tecta** | Temporal Lobe |
| **Piriform Postpiriform Transition Area Combined** | Temporal Lobe |
| **Nucleus of lateral olfactory tract Cortical amygdalar area combined** | Temporal Lobe |
| **Field CA1** | Hippocampal region |
| **Field CA2** | Hippocampal region |
| **Field CA3** | Hippocampal region |
| **Dentate gyrus** | Hippocampal region |
| **Entorhinal area** | Retrohippocampal region |

| | |
|---|---|
| **Parasubiculum** | Retrohippocampal region |
| **Postsubiculum** | Retrohippocampal region |
| **Presubiculum** | Retrohippocampal region |
| **Subiculum** | Retrohippocampal region |
| **Caudoputamen** | Striatum |
| **Nucleus accumbens** | Striatum |
| **Fundus of striatum** | Striatum |
| **Olfactory tubercle** | Striatum |
| **Lateral septal complex** | Striatum |
| **Striatum-like amygdalar nuclei** | Striatum |
| **Globus pallidus, external segment** | Pallidum |
| **Substantia innominata** | Pallidum |
| **Medial septal complex** | Pallidum |
| **Bed nuclei of the stria terminalis** | Pallidum |
| **Ventral group of the dorsal thalamus** | Thalamus |
| **Medial geniculate complex** | Thalamus |
| **Dorsal part of the lateral geniculate complex** | Thalamus |
| **Lateral group of the dorsal thalamus** | Thalamus |
| **Anterior group of the dorsal thalamus** | Thalamus |
| **Medial group of the dorsal thalamus** | Thalamus |
| **Midline group of the dorsal thalamus** | Thalamus |
| **Reticular nucleus of the thalamus** | Thalamus |
| **Periventricular zone** | Hypothalamus |
| **Hypothalamic medial zone** | Hypothalamus |
| **Hypothalamic lateral zone** | Hypothalamus |
| **Zona incerta** | Hypothalamus |

**Supplementary Table 3:** P Values across different time points when quantifying the difference between 5xFAD vs WT functional connectivity values for Hippocampal and Retrohippocampal ROI related functional pairs. All ROI pairs included if pVal < 0.05

| Time point | ROI Pair | p Value for Difference | 5xFAD coefficient (z value) | WT coefficeint (z value) |
|---|---|---|---|---|
| **Month 1.5** | Zona incerta_Parasubiculum | 0.000198586 | -0.030393389 | 0.03495689 |
| **Month 1.5** | Postsubiculum_Field CA2 | 0.002190965 | -0.002037451 | 0.059429371 |
| **Month 1.5** | Parasubiculum_Entorhinal area | 0.012070849 | 0.240930841 | 0.319072296 |
| **Month 1.5** | Postsubiculum_Prelimbic area | 0.020397987 | 0.074636182 | 0.032706723 |
| **Month 1.5** | Parasubiculum_Field CA1 | 0.029550322 | 0.066534871 | 0.132398149 |
| **Month 1.5** | Entorhinal area_Field CA1 | 0.035679175 | 0.212902843 | 0.317068182 |
| **Month 1.5** | Postsubiculum_Orbital area | 0.03780941 | 0.076398416 | 0.036465124 |
| **Month 1.5** | Substantia innominata_Subiculum | 0.038590064 | 0.021445043 | -0.028097156 |
| **Month 1.5** | Parasubiculum_Visual areas | 0.039342986 | -0.021450399 | 0.045657396 |
| **Month 1.5** | Subiculum_Field CA3 | 0.042179584 | 0.181954172 | 0.24597719 |
| **Month 1.5** | Anterior group of the dorsal thalamus_Parasubiculum | 0.043074134 | 0.032458458 | 0.074917875 |

| | | | | |
|---|---|---|---|---|
| **Month 1.5** | Field CA1_Temporal association areas | 0.048677646 | 0.221409475 | 0.333991639 |
| **Month 1.5** | Subiculum_Temporal association areas | 0.049564972 | 0.112082883 | 0.205425338 |
| **Month 2** | Parasubiculum_Auditory areas | 0.00227307 | -0.017112579 | 0.043375871 |
| **Month 2** | Hypothalamic medial zone_Field CA1 | 0.003587923 | 0.006096266 | -0.055225346 |
| **Month 2** | Presubiculum_Field CA2 | 0.006620088 | -0.007557972 | 0.040664093 |
| **Month 2** | Field CA1_Infralimbic area | 0.011125833 | -0.011100297 | 0.031259645 |
| **Month 2** | Reticular nucleus of the thalamus_Field CA2 | 0.013208657 | -0.023688083 | 0.01812921 |
| **Month 2** | Field CA3_Supplemental somatosensory area | 0.014593589 | 0.039776495 | -0.00783931 |
| **Month 2** | Subiculum_Prelimbic area | 0.01525084 | 0.011785923 | 0.075346832 |
| **Month 2** | Subiculum_Infralimbic area | 0.015720544 | 0.016747395 | 0.075787245 |
| **Month 2** | Presubiculum_Temporal association areas | 0.015722331 | -0.002584058 | 0.068529772 |
| **Month 2** | Zona incerta_Postsubiculum | 0.017628469 | 0.036214324 | -0.003097021 |
| **Month 2** | Periventricular zone_Subiculum | 0.018328081 | 0.010236402 | -0.05251105 |
| **Month 2** | Presubiculum_Auditory areas | 0.019793922 | -0.029526943 | 0.025473548 |
| **Month 2** | Periventricular zone_Presubiculum | 0.020321536 | 0.006790981 | -0.038349367 |

| | | | | |
|---|---|---|---|---|
| **Month 2** | Dorsal part of the lateral geniculate complex_Presubiculum | 0.020527578 | 0.06199892 | 0.0227064 |
| **Month 2** | Subiculum_Retrosplenial area | 0.021967016 | 0.083910857 | 0.158275327 |
| **Month 2** | Field CA1_Piriform Postpiriform Transition Area Combined | 0.025970241 | 0.108705782 | 0.041607314 |
| **Month 2** | Parasubiculum_Temporal association areas | 0.028213848 | -0.027432382 | 0.035491191 |
| **Month 2** | Presubiculum_Posterior parietal association areas | 0.032475909 | -0.004267675 | 0.043086509 |
| **Month 2** | Dorsal part of the lateral geniculate complex_Field CA3 | 0.033626744 | 0.256131331 | 0.213429113 |
| **Month 2** | Dentate gyrus_Field CA2 | 0.034592512 | 0.106875172 | 0.146766535 |
| **Month 2** | Field CA1_Agranular insular area | 0.034776912 | 0.089099358 | 0.023748309 |
| **Month 2** | Field CA3_Anterior cingulate area | 0.036509002 | 0.048298151 | 0.092352349 |
| **Month 2** | Bed nuclei of the stria terminalis_Field CA1 | 0.037560237 | -0.038661358 | 0.004111837 |
| **Month 2** | Periventricular zone_Field CA1 | 0.039536398 | -0.011375928 | -0.061371393 |
| **Month 2** | Nucleus accumbens_Field CA2 | 0.039649869 | -0.010050398 | 0.026232816 |
| **Month 2** | Parasubiculum_Nucleus of lateral olfactory tract Cortical amygdalar area combined | 0.042783365 | 0.033257465 | 0.080240812 |
| **Month 2** | Presubiculum_Orbital area | 0.043525524 | 0.045389798 | 0.087925729 |
| **Month 2** | Periventricular zone_Postsubiculum | 0.045520713 | 0.029830016 | -0.013961495 |
| **Month 2** | Field CA1_Supplemental somatosensory area | 0.049838714 | 0.053464262 | 0.011928486 |

| | | | | |
|---|---|---|---|---|
| **Month 4** | Subiculum_Entorhinal area | 0.001979914 | 0.348736597 | 0.455332445 |
| **Month 4** | Presubiculum_Field CA2 | 0.00451735 | 0.022775918 | 0.073320767 |
| **Month 4** | Fundus of striatum_Field CA3 | 0.013376789 | -0.004014788 | 0.033405823 |
| **Month 4** | Presubiculum_Field CA1 | 0.015120068 | 0.086305786 | 0.154701075 |
| **Month 4** | Parasubiculum_Primary somatosensory area | 0.018265561 | 0.162311618 | 0.113455481 |
| **Month 4** | Parasubiculum_Visual areas | 0.019079415 | 0.022416395 | 0.070988453 |
| **Month 4** | Nucleus accumbens_Postsubiculum | 0.02335086 | 0.004768731 | 0.045167933 |
| **Month 4** | Hypothalamic lateral zone_Dentate gyrus | 0.024659749 | 0.015665642 | -0.02415105 |
| **Month 4** | Field CA3_Field CA2 | 0.028936607 | 0.21554313 | 0.269071777 |
| **Month 4** | Field CA2_Visceral area | 0.034267094 | 0.010301424 | -0.021734895 |
| **Month 4** | Presubiculum_Dentate gyrus | 0.040825263 | 0.301129077 | 0.376468931 |
| **Month 4** | Subiculum_Dentate gyrus | 0.040899698 | 0.346947939 | 0.405713526 |
| **Month 4** | Subiculum_Presubiculum | 0.040915105 | 0.230825791 | 0.294028741 |
| **Month 4** | Subiculum_Parasubiculum | 0.041036495 | 0.191324126 | 0.257849754 |
| **Month 4** | Presubiculum_Secondary motor area | 0.041706645 | 0.056452089 | 0.101874311 |
| **Month 4** | Parasubiculum_Supplemental somatosensory area | 0.044909823 | 0.090471761 | 0.058083831 |
| **Month 4** | Presubiculum_Entorhinal area | 0.046070705 | 0.142795507 | 0.20304892 |

| | | | | |
|---|---|---|---|---|
| **Month 6** | Dentate gyrus_Field CA2 | 3.35251E-06 | 0.071166053 | 0.1596494 |
| **Month 6** | Subiculum_Secondary motor area | 1.10941E-05 | 0.018162662 | 0.113522656 |
| **Month 6** | Postsubiculum_Field CA3 | 5.06715E-05 | -0.011150777 | 0.076707025 |
| **Month 6** | Postsubiculum_Entorhinal area | 8.27713E-05 | 0.038793433 | 0.127547018 |
| **Month 6** | Presubiculum_Entorhinal area | 0.000162935 | 0.095958396 | 0.195892621 |
| **Month 6** | Subiculum_Field CA2 | 0.000193048 | -0.004160883 | 0.059580541 |
| **Month 6** | Postsubiculum_Field CA1 | 0.000375432 | 0.007906686 | 0.088410988 |
| **Month 6** | Subiculum_Anterior cingulate area | 0.000629217 | 0.014666639 | 0.093012965 |
| **Month 6** | Anterior group of the dorsal thalamus_Presubiculum | 0.001084376 | 0.042231518 | 0.105575495 |
| **Month 6** | Lateral group of the dorsal thalamus_Subiculum | 0.001562017 | 0.054652046 | 0.120852937 |
| **Month 6** | Presubiculum_Field CA1 | 0.002094266 | 0.048904262 | 0.127279758 |
| **Month 6** | Zona incerta_Postsubiculum | 0.002555657 | -0.012111784 | 0.045517989 |
| **Month 6** | Field CA3_Secondary motor area | 0.004218065 | 0.010221668 | 0.066327716 |
| **Month 6** | Medial group of the dorsal thalamus_Field CA3 | 0.004427622 | 0.009149334 | 0.056864034 |
| **Month 6** | Lateral group of the dorsal thalamus_Dentate gyrus | 0.004692743 | 0.191410431 | 0.265645858 |
| **Month 6** | Field CA1_Primary motor area | 0.004925751 | -0.016176565 | 0.041746437 |
| **Month 6** | Subiculum_Primary motor area | 0.005338795 | 0.026607688 | 0.096044354 |

| | | | | |
|---|---|---|---|---|
| **Month 6** | Postsubiculum_Nucleus of lateral olfactory tract Cortical amygdalar area combined | 0.005840674 | -0.005514111 | 0.058820208 |
| **Month 6** | Medial geniculate complex_Entorhinal area | 0.005863774 | 0.031065908 | 0.07624775 |
| **Month 6** | Anterior group of the dorsal thalamus_Subiculum | 0.006082382 | 0.024573284 | 0.081796365 |
| **Month 6** | Reticular nucleus of the thalamus_Field CA1 | 0.006468457 | -0.046078966 | 0.001018094 |
| **Month 6** | Entorhinal area_Anterior cingulate area | 0.00647736 | 0.018191046 | 0.079948139 |
| **Month 6** | Postsubiculum_Piriform Postpiriform Transition Area Combined | 0.007674961 | -0.00198748 | 0.073573815 |
| **Month 6** | Ventral group of the dorsal thalamus_Presubiculum | 0.008125318 | 0.014111355 | 0.067251044 |
| **Month 6** | Globus pallidus, external segment_Postsubiculum | 0.008191446 | 0.005520778 | -0.033255075 |
| **Month 6** | Parasubiculum_Infralimbic area | 0.009576736 | 0.030039053 | 0.07088126 |
| **Month 6** | Lateral group of the dorsal thalamus_Presubiculum | 0.009644414 | 0.068391734 | 0.12920925 |
| **Month 6** | Anterior group of the dorsal thalamus_Postsubiculum | 0.010595782 | 0.030690354 | 0.088998761 |
| **Month 6** | Anterior group of the dorsal thalamus_Dentate gyrus | 0.010814894 | 0.119464299 | 0.179054832 |
| **Month 6** | Field CA2_Retrosplenial area | 0.011139452 | 0.010981447 | 0.055103234 |
| **Month 6** | Dentate gyrus_Field CA3 | 0.012188611 | 0.348603395 | 0.412241496 |
| **Month 6** | Hypothalamic lateral zone_Postsubiculum | 0.012531789 | -0.024731295 | 0.030424615 |
| **Month 6** | Subiculum_Field CA3 | 0.013731003 | 0.10488591 | 0.173645437 |
| **Month 6** | Field CA1_Orbital area | 0.013759529 | 0.00137189 | 0.047018017 |

| | | | | |
|---|---|---|---|---|
| **Month 6** | Presubiculum_Orbital area | 0.013841129 | 0.030271482 | 0.079604361 |
| **Month 6** | Field CA3_Prelimbic area | 0.014261991 | -0.006809092 | 0.036029149 |
| **Month 6** | Presubiculum_Field CA2 | 0.017781282 | 0.003854877 | 0.048931144 |
| **Month 6** | Field CA3_Anterior cingulate area | 0.018200363 | 0.016666347 | 0.060545474 |
| **Month 6** | Dentate gyrus_Secondary motor area | 0.018635921 | 0.091310537 | 0.155296894 |
| **Month 6** | Medial geniculate complex_Field CA2 | 0.019026236 | -0.012375085 | 0.024495737 |
| **Month 6** | Caudoputamen_Entorhinal area | 0.019493586 | 0.028643455 | 0.086738172 |
| **Month 6** | Entorhinal area_Secondary motor area | 0.019593459 | 0.094503653 | 0.158150235 |
| **Month 6** | Presubiculum_Piriform Postpiriform Transition Area Combined | 0.02047872 | 0.015382854 | 0.066326783 |
| **Month 6** | Midline group of the dorsal thalamus_Presubiculum | 0.022391254 | 0.018402106 | 0.057383036 |
| **Month 6** | Field CA1_Anterior cingulate area | 0.022489192 | 0.003801188 | 0.055385185 |
| **Month 6** | Dentate gyrus_Anterior cingulate area | 0.022929871 | 0.083593743 | 0.149892207 |
| **Month 6** | Postsubiculum_Taenia tecta | 0.023031474 | 0.058175324 | 0.005041997 |
| **Month 6** | Presubiculum_Infralimbic area | 0.026505797 | 0.047600083 | 0.08436003 |
| **Month 6** | Entorhinal area_Infralimbic area | 0.027508882 | 0.021749997 | 0.07799372 |
| **Month 6** | Medial geniculate complex_Field CA1 | 0.032003374 | 0.033156488 | 0.077050793 |
| **Month 6** | Reticular nucleus of the thalamus_Presubiculum | 0.034425105 | -0.008278541 | 0.030417404 |
| **Month 6** | Reticular nucleus of the thalamus_Subiculum | 0.035206295 | 0.000960418 | 0.04097793 |

| | | | | |
|---|---|---|---|---|
| **Month 6** | Dorsal part of the lateral geniculate complex_Subiculum | 0.035819221 | 0.03633991 | 0.085377164 |
| **Month 6** | Field CA1_Prelimbic area | 0.037390499 | -0.00490207 | 0.037395559 |
| **Month 6** | Caudoputamen_Field CA2 | 0.037897227 | 0.021015911 | -0.014220356 |
| **Month 6** | Ventral group of the dorsal thalamus_Field CA2 | 0.038447528 | 0.002839833 | 0.034108561 |
| **Month 6** | Presubiculum_Prelimbic area | 0.039038011 | 0.035039073 | 0.082108686 |
| **Month 6** | Midline group of the dorsal thalamus_Field CA1 | 0.039329464 | 0.005998328 | 0.045427617 |
| **Month 6** | Parasubiculum_Entorhinal area | 0.040902768 | 0.176681635 | 0.22960052 |
| **Month 6** | Postsubiculum_Secondary motor area | 0.042695946 | 0.046564203 | 0.094344274 |
| **Month 6** | Presubiculum_Field CA3 | 0.043046799 | 0.047195086 | 0.101860866 |
| **Month 6** | Zona incerta_Dentate gyrus | 0.043837529 | 0.028484108 | 0.071413011 |
| **Month 6** | Parasubiculum_Anterior cingulate area | 0.04388488 | 0.035619277 | 0.078743268 |
| **Month 6** | Fundus of striatum_Dentate gyrus | 0.046176303 | 0.005627813 | 0.044855985 |
| **Month 6** | Entorhinal area_Primary motor area | 0.04838384 | 0.104524939 | 0.160362121 |
| **Month 6** | Medial group of the dorsal thalamus_Dentate gyrus | 0.049544419 | 0.082486926 | 0.12563569 |

**Supplementary Table 4**: Functional Connectivity values that were deemed significant under p < 0.005

| Timepoint | ROI Pair | p Value for Difference | 5xFAD coefficient (z value) | WT coefficeint (z value) |
|---|---|---|---|---|
| Month 1.5 | Zona incerta_Parasubiculum | 0.000198586 | -0.030393389 | 0.03495689 |
| Month 1.5 | Dorsal part of the lateral geniculate complex_Temporal association areas | 0.001105172 | 0.02946571 | 0.094627917 |
| Month 1.5 | Postsubiculum_Field CA2 | 0.002190965 | -0.002037451 | 0.059429371 |
| Month 2 | Taenia tecta_Primary somatosensory area, nose | 0.000131648 | 0.057590928 | -0.046631038 |
| Month 2 | Caudoputamen_Primary somatosensory area, nose | 0.001289112 | 0.237982583 | 0.121855404 |
| Month 2 | Striatum-like amygdalar nuclei_Prelimbic area | 0.001544699 | 0.038611229 | -0.048601923 |
| Month 2 | Parasubiculum_Auditory areas | 0.00227307 | -0.017112579 | 0.043375871 |
| Month 2 | Medial group of the dorsal thalamus_Primary somatosensory area, nose | 0.002904204 | 0.067968378 | 0.005308862 |
| Month 2 | Fundus of striatum_Primary somatosensory area, nose | 0.003532272 | 0.078812834 | 0.015817962 |
| Month 2 | Hypothalamic medial zone_Field CA1 | 0.003587923 | 0.006096266 | -0.055225346 |
| Month 2 | Posterior parietal association areas_Visual areas | 0.004032598 | 0.160938965 | 0.269131805 |
| Month 2 | Medial septal complex_Striatum-like amygdalar nuclei | 0.004290314 | 0.061478933 | -0.025276181 |
| Month 2 | Midline group of the dorsal thalamus_Bed nuclei of the stria terminalis | 0.004766883 | 0.139312078 | 0.078262734 |

| | | | | |
|---|---|---|---|---|
| Month 4 | Anterior group of the dorsal thalamus_Striatum-like amygdalar nuclei | 8.60422E-06 | 0.027894588 | -0.061234621 |
| Month 4 | Ectorhinal area_Temporal association areas | 0.000400764 | 0.557915032 | 0.7224864 |
| Month 4 | Globus pallidus, external segment_Fundus of striatum | 0.001509045 | 0.077070442 | 0.016939897 |
| Month 4 | Subiculum_Entorhinal area | 0.001979914 | 0.348736597 | 0.455332445 |
| Month 4 | Lateral septal complex_Visceral area | 0.003111034 | 0.026869985 | -0.041681804 |
| Month 4 | Presubiculum_Field CA2 | 0.00451735 | 0.022775918 | 0.073320767 |
| Month 6 | Dentate gyrus_Field CA2 | 3.35251E-06 | 0.071166053 | 0.1596494 |
| Month 6 | Subiculum_Secondary motor area | 1.10941E-05 | 0.018162662 | 0.113522656 |
| Month 6 | Postsubiculum_Field CA3 | 5.06715E-05 | -0.011150777 | 0.076707025 |
| Month 6 | Postsubiculum_Entorhinal area | 8.27713E-05 | 0.038793433 | 0.127547018 |
| Month 6 | Presubiculum_Entorhinal area | 0.000162935 | 0.095958396 | 0.195892621 |
| Month 6 | Subiculum_Field CA2 | 0.000193048 | -0.004160883 | 0.059580541 |
| Month 6 | Postsubiculum_Field CA1 | 0.000375432 | 0.007906686 | 0.088410988 |
| Month 6 | Subiculum_Anterior cingulate area | 0.000629217 | 0.014666639 | 0.093012965 |
| Month 6 | Anterior group of the dorsal thalamus_Gustatory areas | 0.000693186 | -0.036410685 | 0.040784194 |
| Month 6 | Anterior group of the dorsal thalamus_Presubiculum | 0.001084376 | 0.042231518 | 0.105575495 |
| Month 6 | Lateral septal complex_Piriform Postpiriform Transition Area Combined | 0.001288713 | -0.075652057 | 0.023271878 |
| Month 6 | Lateral group of the dorsal thalamus_Subiculum | 0.001562017 | 0.054652046 | 0.120852937 |
| Month 6 | Bed nuclei of the stria terminalis_Fundus of striatum | 0.001910656 | -0.017999218 | 0.052353521 |

| | | | | |
|---|---|---|---|---|
| Month 6 | Dorsal part of the lateral geniculate complex_Auditory areas | 0.002069057 | -0.004777761 | 0.054111352 |
| Month 6 | Presubiculum_Field CA1 | 0.002094266 | 0.048904262 | 0.127279758 |
| Month 6 | Reticular nucleus of the thalamus_Ventral group of the dorsal thalamus | 0.002330891 | 0.25530997 | 0.324207618 |
| Month 6 | Zona incerta_Postsubiculum | 0.002555657 | -0.012111784 | 0.045517989 |
| Month 6 | Midline group of the dorsal thalamus_Visceral area | 0.002626394 | -0.011883269 | 0.039551044 |
| Month 6 | Zona incerta_Striatum-like amygdalar nuclei | 0.002649032 | 0.003572915 | 0.106442471 |
| Month 6 | Lateral septal complex_Auditory areas | 0.002916453 | -0.016645682 | 0.03591924 |
| Month 6 | Bed nuclei of the stria terminalis_Ectorhinal area | 0.003048788 | -0.036717491 | 0.012314602 |
| Month 6 | Ventral group of the dorsal thalamus_Gustatory areas | 0.003080705 | -0.015412888 | 0.045122949 |
| Month 6 | Taenia tecta_Secondary motor area | 0.003485417 | 0.15127597 | 0.027819241 |
| Month 6 | Medial septal complex_Piriform Postpiriform Transition Area Combined | 0.003500702 | -0.115103318 | -0.025987267 |
| Month 6 | Field CA3_Secondary motor area | 0.004218065 | 0.010221668 | 0.066327716 |
| Month 6 | Medial group of the dorsal thalamus_Field CA3 | 0.004427622 | 0.009149334 | 0.056864034 |
| Month 6 | Lateral group of the dorsal thalamus_Dentate gyrus | 0.004692743 | 0.191410431 | 0.265645858 |
| Month 6 | Field CA1_Primary motor area | 0.004925751 | -0.016176565 | 0.041746437 |
